# Evaluating BOLD functional MRI biophysical simulation approaches: impact of vascular geometry, magnetic field calculations, and water diffusion models

**DOI:** 10.1101/2025.08.29.673098

**Authors:** Avery J. L. Berman, Jacob Chaussé, Grant Hartung, Jonathan R. Polimeni, J. Jean Chen

## Abstract

Biophysical simulations have guided the development of blood oxygenation level-dependent (BOLD) functional MRI (fMRI) acquisitions and signal models that relate the BOLD signal to the underlying physiology, such as calibrated BOLD and vascular fingerprinting. Numerous simulation techniques have been developed, however, few of them have been directly compared, thus limiting the assessment of the accuracy and interchangeability of these methods as well as the accuracy of the quantitative techniques derived from them. In this work, we compared the accuracy and computational demands of eight previously published simulation approaches that adopt different geometries (ranging from infinite cylinders to synthetic vascular anatomical networks (VANs)), field offset calculations (analytical and Fourier-based), and water diffusion implementations (Monte Carlo and convolution-based), all of which are available in an open-source Python toolkit, *BOLDsωimsuite*. The reference simulation approach for comparison used three-dimensional infinite cylinders, analytical field offsets, and Monte Carlo diffusion. When compared with the reference approach, most of the simulations, including two- and three-dimensional geometries, were in excellent agreement when assuming the intravascular signal contribution was small. Two commonly employed simulation approaches were notably biased; both used two-dimensional geometries with overly simplified vasculature or field offset calculations. In general, the simulated intravascular signal was the least consistent across approaches, thus potentially resulting in larger errors when the intravascular signal contribution is large. Lastly, the VAN results were in good agreement with the reference but they diverged slightly, yet systematically, from each other at smaller radii (≲ 3 μm), primarily driven by intravascular signal differences. We conclude, therefore, that the reference approach is an attractive option for exploratory simulations in the many cases where anatomical and hemodynamic realism is not needed, balancing ease of implementation, accessibility, versatility, computational efficiency, accuracy of results, and interpretability. These findings help pave the way for a broader adoption of forward modelling of the BOLD signal and more reliable interpretations of biophysical simulations aiming to develop quantitative models of the BOLD signal.

## 1 Introduction

Ever since its discovery, the blood oxygenation level-dependent (BOLD) MRI signal (Ogawa et al., 1990) has been the workhorse for mapping the neuronal basis of human cognition and behaviour. Upon neuronal activation, a complex combination of vascular and metabolic changes gives rise to local changes in the magnetic susceptibility of blood, ultimately resulting in the observed intensity changes on, typically, *T*_2_*-weighted images (Bandettini et al., 1992; Kwong et al., 1992; Ogawa et al., 1992). The continual development of new hardware (Feinberg et al., 2023) and pulse sequences (Berman, Grissom, et al., 2021; Setsompop et al., 2012) has allowed scientists to reliably map brain-wide changes in neural activity at high spatial (Bollmann & Barth, 2021) and temporal (Polimeni & Lewis, 2021) resolutions using BOLD functional MRI (fMRI). While invasive animal studies have greatly improved our understanding of neurovascular coupling (Iadecola, 2017), interpreting the BOLD signal remains non-trivial for clinical applications (Pike, 2012) and novel experimental paradigms, like those invoking rapid or transient BOLD responses (Polimeni & Lewis, 2021). In this regard, biophysical simulations have been foundational to understanding how physiological parameters, e.g., blood volume or vessel size, and acquisition choices, e.g., spin-echo or gradient-echo sequences, impact BOLD contrast (Bandettini & Wong, 1995; Boxerman, Hamberg, et al., 1995; Gagnon et al., 2015; Martindale et al., 2008; Ogawa et al., 1993; Uludag et al., 2009). Simulations have been used to guide fMRI acquisitions with greater BOLD detection sensitivity and physiological specificity (Baez-Yanez et al., 2017; Boxerman, Hamberg, et al., 1995; Chen & Calhoun, 2012; Miller & Jezzard, 2008; Pflugfelder et al., 2011; Scheffler et al., 2019, 2021), and they have helped shape the development of many techniques for quantitatively imaging brain physiology, including calibrated BOLD (Berman et al., 2018; Davis et al., 1998; Griffeth & Buxton, 2011), quantitative BOLD (Dickson et al., 2010; Stone et al., 2019; Yablonskiy & Haacke, 1994), vascular fingerprinting (Christen et al., 2014; Delphin et al., 2024), and perfusion imaging with dynamic susceptibility contrast (DSC) (Boxerman, Hamberg, et al., 1995). Thus, biophysical simulations may hold the key to unleashing the full potential of fMRI in future applications; however, comparison across simulation approaches is often lacking, and as techniques like calibrated BOLD and vascular fingerprinting are quantitative, it is important to understand how the choice of simulation approach on which they are based influences their results (X. Cheng et al., 2019; Gagnon et al., 2016; Griffeth & Buxton, 2011; Martindale et al., 2008).

Since the early days of fMRI, several simulation approaches have been developed, trading off anatomical and hemodynamic realism, model complexity, and computational demand. Across these approaches, the major variations that have been developed typically relate to one of three major simulation stages: (i) the vascular geometry definition, including random infinite cylinders in three-dimensions (3D) or two-dimensions (2D) (Bandettini & Wong, 1995), or more anatomically realistic vessel networks, composed of finite, branching cylinders, commonly referred to as vascular anatomical networks (VANs) (Gagnon et al., 2015). (ii) The main magnetic field (*B*_0_) offset calculation, including analytically calculated or Fourier-based field maps (Marques & Bowtell, 2008), continuous or discrete field maps, single-vessel techniques simulated and averaged over multiple *B*_0_ angles (Ogawa et al., 1993), or averaging *B*_0_ maps over multiple angles before simulation (Pannetier et al., 2013). And (iii) the implementation of water diffusion, including Monte Carlo (MC) methods, so-called deterministic diffusion, which is a convolution-based spreading of magnetization (Bandettini & Wong, 1995; Doucette et al., 2018), or a finite-difference solution to the Bloch equations with diffusion (Marques & Bowtell, 2008). Most simulation approaches are independently developed for each user’s application; thus, they are rarely directly compared with each other, making it challenging to know whether their results agree or not.

In this study, we implemented and compared the most commonly used simulation approaches from the literature, including the combination of three different vascular geometries (with recent advances using synthetic VANs), six *B*_0_ offset calculation methods, and two water diffusion models, totalling eight unique approaches. To ensure consistency across simulations, all techniques were implemented within a single toolkit (Chaussé et al., 2025). Whereas most of the studies discussed above (e.g.,(Boxerman, Hamberg, et al., 1995; Martindale et al., 2008; Uludag et al., 2009), etc.) examined the impact of a range of simulation parameter values—such as net field offset or blood volume—on transverse signal decay using their preferred simulation approach, we confined our simulations to a limited range of parameter values and, instead, evaluated the effect of the simulation approaches themselves. Since analytically defined solutions for the simulations from VAN models are not yet available, we evaluated the accuracy of the simulations relative to a reference technique based on 3D Monte Carlo simulations with random infinite cylinders. Infinite cylinders are commonly used in BOLD signal simulations: they possess analytical solutions to their *B*_0_ offset calculations and their resulting signal in some instances, they were the first geometry used in seminal studies to derive key predictions about the signal from endogenous and exogenous sources—i.e., deoxyhemoglobin and contrast agents, respectively—of *T*_2_ and *T*_2_* contrast, and these simulations have been experimentally verified (Yablonskiy, 1998). Many of the simplified approaches that were later developed were derived from infinite cylinders, so the comparison to infinite cylinders as a reference naturally makes sense. From this validated reference, we then incrementally changed our simulation approaches such that we could isolate the effect of the simulation design choices (e.g., the effect of *B*_0_ offset calculation from vascular geometry). The following Guiding Questions informed our experiments and their interpretation:

#1. For a given application, how does the simulation approach affect the outcomes?
#2. How similar are the results of the infinite cylinder simulation approaches, and what are their key differences?
#3. How do the results of simulations from infinite cylinders and simplified VANs differ, and what drives these differences? I.e., what is the effect of assuming blood vessels are infinite cylinders?

For Guiding Question #1, we used a small-scale vascular fingerprinting analysis and examined the accuracy of the estimated vessel radii for the various simulation approaches. This fingerprinting experiment and the remaining analyses, which directly compared simulation accuracy, helped address Guiding Questions #2 and #3. Although most approaches demonstrated good agreement with the reference approach, the oversimplification in some commonly used methods resulted in high levels of error. Furthermore, our results demonstrate that most 2D and 3D simulations using infinite cylinders provide a good approximation of simulations from more realistic capillary beds, but they may need to be adapted to capture the fMRI signal behaviour across the full microvascular hierarchy, which contains more structured and asymmetric vessel orientations (i.e., intra-cortical and pial vessels) and a range of blood oxygenation values exhibiting a rough spatial organization imparted by the connections from arteries to capillaries to veins.

## 2 Theory

All simulations can be broken down into four stages: (1) defining the geometry, where the geometry of the simulation voxel and the vessels is determined; (2) calculating the *B*_0_ field offsets; (3) modelling the self-diffusion of water molecules; and (4) generating the MR signal for a given pulse sequence. In all simulations, we calculate the signal resulting from the relative dephasing of transverse magnetization from vessel-induced field inhomogeneities. Longitudinal magnetization and intrinsic *T*_1_ and *T*_2_ relaxation were ignored in this study. In the following, we review the theory that underlies all the simulation approaches that we tested, starting from stage (2), the field offset calculations.

### 2.1 Field offset calculations

The *B*_0_ field offsets, Δ*B*_0_, generated by partially deoxygenated blood in vessels are generally present in the intravascular (IV) space and the surrounding extravascular (EV) space. A common approach is to represent the vessels by infinitely long cylinders since Δ*B*_0_ can then be calculated analytically. For an infinite cylinder of radius *R*, the field offsets are given by (Ogawa et al., 1993):

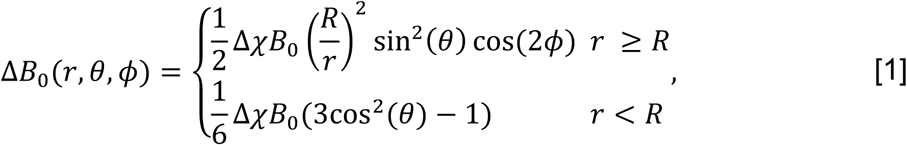

where Δ*χ* is the susceptibility difference between the blood within the vessel and the surrounding tissue, *r* is the Euclidean distance from the point of interest to the vessel axis, *θ* is the angle between *B*_0_ and the vessel axis, and *ϕ* is the angle between the projection of *B*_0_ onto the plane perpendicular to the vessel axis and the line segment connecting the point of interest and the vessel axis. Fig. 1A and 1B depict the vessel geometry relative to *B*_0_ and the corresponding Δ*B*_0_ map, respectively.

**Fig. 1:**
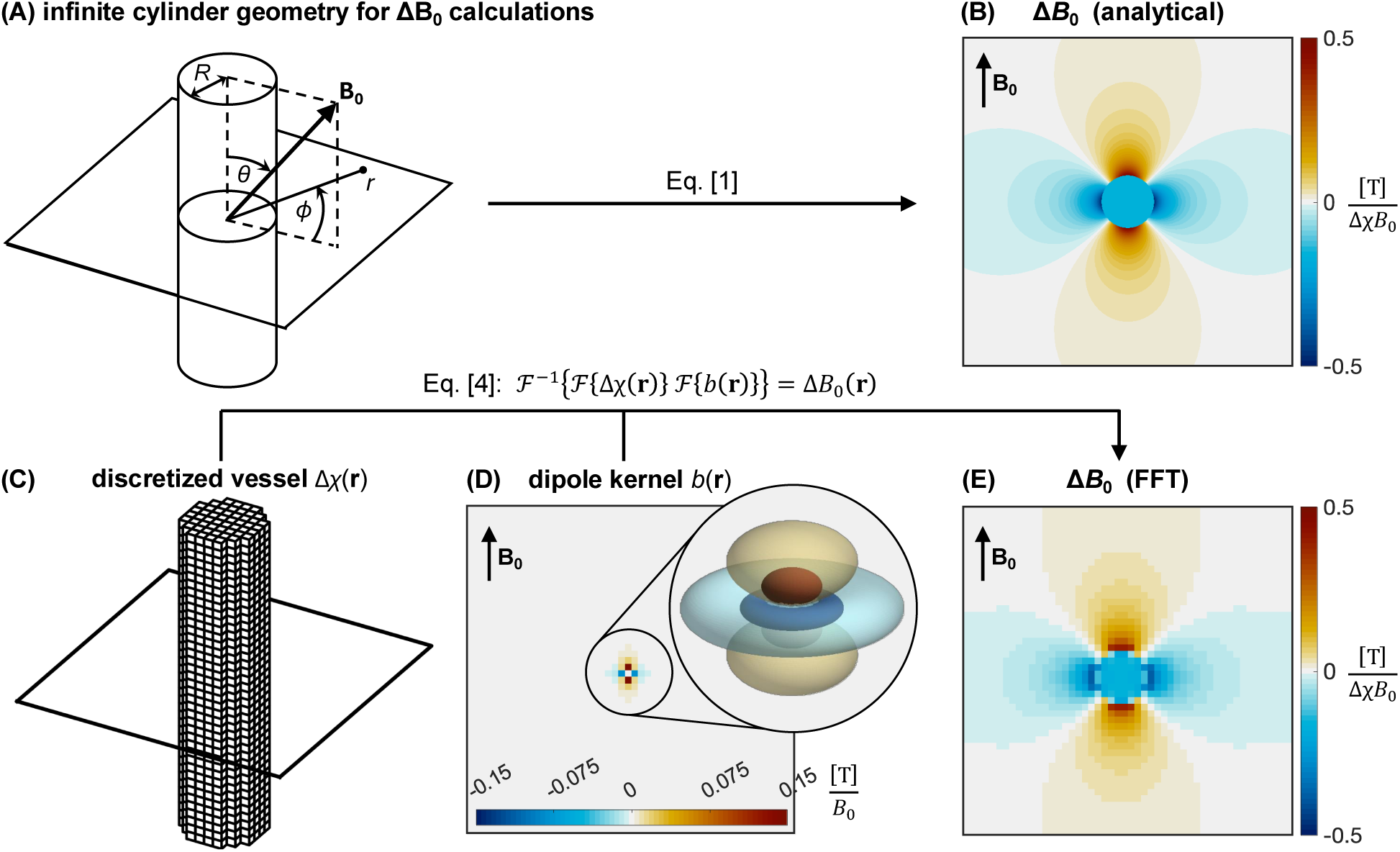
(A) The geometry used in Eq. [1] for determining the field offset a distance *r* away from the axis of an infinite cylinder of radius *R* in an external field, **B_0_**. (B) A cross-section of the analytically calculated field offsets produced by an infinite cylinder of uniform magnetic susceptibility, based on Eq. [1]. (C–D) Components of the Fourier-based Δ*B*_0_ calculation based on Eq. [4], including (C) the discretized susceptibility distribution of an infinite cylinder (i.e., blood vessel), (D) the dipole kernel, and (E), the resulting discretized Δ*B*_0_ map. In (D), the dipole kernel is the field perturbation from a single voxel and is, thus, of coarse resolution and difficult to visualize in a two-dimensional slice. The inset shows a three-dimensional rendering of how the kernel would appear if it could be resolved at a higher spatial resolution, which corresponds to the field of a sphere.

For an arbitrary spatial distribution of magnetic susceptibility, such as a realistic representation of the vasculature with branching, tortuous vessels (Baez-Yanez et al., 2017; Gagnon et al., 2015; Hartung et al., 2018), Δ*B*_0_ can be calculated by a forward model where the spatially varying magnetic susceptibility, Δ*χ*(**r**), is convolved with the field offsets produced by a point-source dipole, *b*(**r**), (Marques & Bowtell, 2005; Salomir et al., 2003):

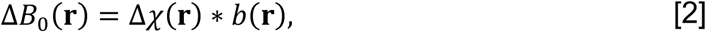

where ∗ is the convolution operator. When *B*_0_ is along the *z*-direction, the point-source dipole field is given by

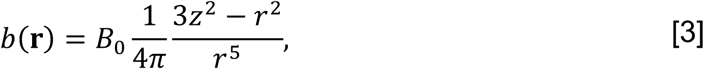

where *r* is the distance from the dipole centre to the point of interest, and *z* is the longitudinal distance from the dipole centre to the point of interest. The efficiency of the convolution can be improved by performing a multiplication in *k* space by means of the Fourier transform (Y. C. Cheng et al., 2009):

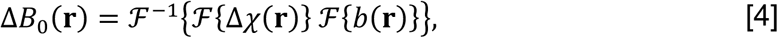

where ℱ and ℱ^-1^ represent the forward and inverse three-dimensional Fourier transforms, respectively. For most geometries, the Fourier transform must be calculated numerically using discrete data, therefore, Δ*B*_0_ will have a spatial resolution that is limited by the discretization of Δ*χ*(**r**). In contrast, when Δ*B*_0_ is calculated for infinite cylinders using the analytic expression in Eq. [1], the spatial coordinates can be continuously defined within machine precision.

### 2.2 Diffusion and dephasing calculations

The self-diffusion of water plays a significant role in MRI, resulting in a vessel-size dependence of BOLD contrast (Boxerman, Hamberg, et al., 1995). For MRI, there are two common approaches to model the diffusion of water that were tested here: the Monte Carlo method and the deterministic diffusion method. In all cases, we considered independent and isotropic diffusion along all dimensions with a diffusion coefficient, *D*.

#### 2.2.1 Monte Carlo diffusion

In the MC method, individual spin-bearing particles independently follow a random walk. At each time point, the position of each spin is displaced by a random amount along each dimension drawn from a Gaussian distribution with a mean of zero and variance *σ*^2^ = 2*Dδt*, where *δt* is the timestep. If we model the diffusion of *N_s_* particles over *N_t_* time points, the position of each particle over time can be represented by the vector **r***_j,k_*, where *j* indexes time and *k* indexes particles. As the particles pass through the field offsets generated by blood vessels, their spins accrue a net phase

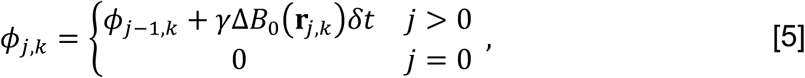

where *γ* = 2.675×10^8^ rad⋅s^−1^⋅T^−1^ is the gyromagnetic ratio for hydrogen nuclei, and Δ*B*_0_(**r***_j,k_*) is the net field offset from all vessels at the position **r***_j,k_*. Immediately after excitation, all spins have 0 phase, corresponding to real-valued magnetization. To model a spin-echo signal at time point *T_E_*, a refocusing pulse was modelled as an instantaneous 180° rotation of the spins about the real axis at time point *T*_180_ = *T_E_*/*2*, resulting in *ϕ*_*T*_180_,*k*_ = –*ϕ*_*T*_180_-1,*k*_.

Finally, the EV, IV, and total signals at the *j*-th time step are given by:

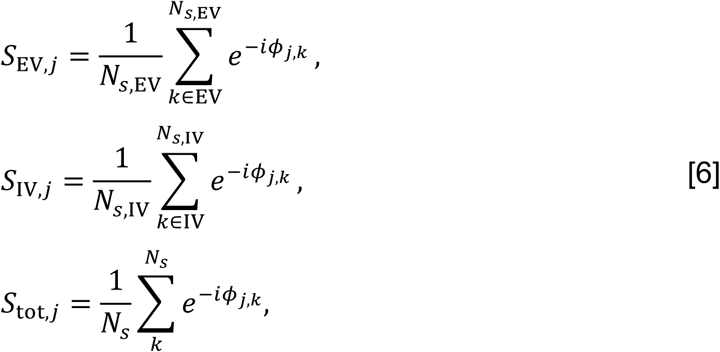

where *N_s,X_* (*X* ∈ {EV, IV}) are the number of spins that are in the EV or IV space, and the sums are only over those spins which reside in their respective spaces.

#### 2.2.2 Deterministic diffusion

In deterministic diffusion (DD), rather than track the spin of individual particles, the transverse magnetization across the voxel is discretized on a grid (Bandettini & Wong, 1995). Solving the diffusion equation by the method of Green’s functions, the diffusion of spins can be represented by the convolution of the magnetization with a Gaussian diffusion kernel, resulting in the progressive blurring of the magnetization over time. If we represent the magnetization of the voxel as a complex-valued matrix, **M**, the resulting magnetization at the *j*-th timepoint after diffusion and precession can be expressed as (Bandettini & Wong, 1995; Doucette et al., 2018)

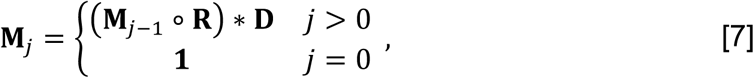

where ∘ is the element-wise multiplication operator and ∗ is still the convolution operator. **M**_0_ = **1** implies that the initial magnetization after excitation is real-valued and set to 1 at all grid points, assuming uniform proton density and ideal excitation. **R** captures the precession at each grid point due to the net field offset, i.e.,

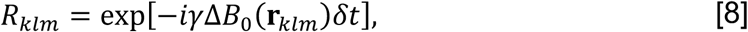

where *k*, *l*, and *m* represent indices in three-dimensional space and Δ*B*_0_ can be calculated as described above in section 2.1. **D** is the diffusion matrix, representing the average displacement of all spin particles within a grid element per unit time. For independent diffusion along each dimension, **D** can be defined in one dimension using the solution to the discretized diffusion equation (Lindeberg, 1990),

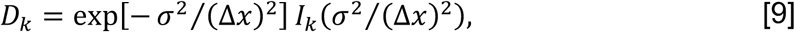

where *k* is the grid index (centred on 0), *σ*^2^ is the variance defined above, Δ*x* is the grid element spacing, and *I_k_* is the modified Bessel function of the first kind. When the diffusion kernel is highly sampled, it approaches the continuous Gaussian distribution as desired (Pannetier et al., 2014). To model a spin-echo signal at time index *T_E_*, a refocusing pulse was modelled as an instantaneous 180° rotation of the magnetization at each grid point about the real axis at time index *T*_180_ = *T_E_*/2, resulting in element-wise complex conjugation of **M**.

The total magnetization can be separated into its EV and IV contributions

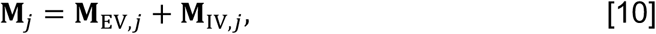

and the EV, IV, and total signals at the *j*-th time step are given by

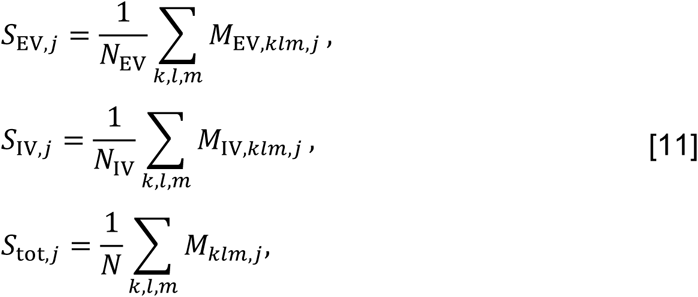

where *N* is the number of grid elements in the voxel, and *N_X_* (*X* ∈ {EV, IV}) is the number of EV or IV grid elements.

## 3 Methods

### 3.1 Signal simulations

All simulations were performed in Python v3.10 using an in-house developed toolkit, *BOLDSωimSuite* (https://github.com/jacobchausse/BOLDswimsuite) (Chaussé et al., 2025). For all simulation techniques (described below), the simulation parameters and their values are summarized in Table 1, and additional details are provided below. All simulations ran single-threaded on a 136-node computing cluster with 187 GB of memory per node, 44 cores per node, and Intel Xeon Gold 6238 2.10-GHz CPUs.

**Table 1:** The common simulation parameter values used. MC: Monte Carlo; TE: echo-time.

| Parameter | Value |
| --- | --- |
| number of voxels | 10 |
| blood volume fraction | 2% |
| vessel radii ($\mu\text{m}$) | 1, 1.5, 2, 3, 4, 5, 6, 8, 10, 13, 16, 20, 30, 40, 60 |
| vessel permeability | 0 |
| $B_0$ | 3 T |
| susceptibility offset, $\Delta\chi$ | $4.15 \times 10^{-7}$ (SI units) |
| diffusion coefficient, $D$ | $1 \mu\text{m}^2/\text{ms}$ |
| # spins per voxel (MC methods) | 40,000 |
| spin-echo TE | 70 ms |
| gradient-echo TE | 30 ms |
| timestep, $\delta t$ | 0.2 ms |
| signal simulation time | 120 ms |

#### 3.1.1 Voxel/geometry definition

Ten voxels were randomly populated with 1-μm radius vessels to a volume fraction of 2%, reflecting predominantly cortical blood volume of deoxygenated blood vessels, i.e., capillaries and veins (Blockley et al., 2013). Note that 1 μm is slightly smaller than a typical capillary radius (Blinder et al., 2013; Hartung, Badr, Mihelic, et al., 2021; Lauwers et al., 2008), but was used here to more thoroughly characterize diffusion effects in the motional narrowing regime, which may be encountered if the diffusion coefficient is elevated. To simulate the signal for larger vessels of radius *R* (up to 60-μm), we reused the original voxels but assigned a side-length scaled by a factor *R*/*R*_0_, where *R*_0_ = 1 μm, as commonly performed (Boxerman, Hamberg, et al., 1995; Martindale et al., 2008; Pannetier et al., 2013; Stone et al., 2019). The voxel size was chosen to be large enough to capture the long-range effects of vessels (Boxerman, Hamberg, et al., 1995): in 3D, voxels had an isotropic edge length of 149 μm, resulting in approximately 400 vessels per voxel for the infinite cylinder approaches; in 2D, an edge length of 251 μm was used to also give 400 vessels per voxel (except for the single-vessel approach, described below in 3.2 Simulation approaches (2D-ANA-MC-1V)). See Supplementary Material Table S1 for the complete listing of effective voxel sizes and grid sizes across methods.

#### 3.1.2 Δ*B*_0_ calculation

*B*_0_ offsets were calculated in continuous space or on a grid. For simulations using a grid, 3D voxels contained 500 grid points per side (6.7 grid points per vessel diameter) and 2D voxels contained at least 1000 grid points per side (8 grid points per vessel diameter). These resolutions were set based on when the root mean square deviation between the reference simulation (see below) and gridded simulations came to a lower plateau (results shown in Supplementary Material Fig. S1). The reason for the increased resolution in 2D is explained below in 3.2 Simulation approaches (2D-ANA-DD).

#### 3.1.3 Diffusion calculation

For MC simulations, a cyclic boundary condition was implemented for the spins, where any spins that reached the edge of the voxel leave the domain on one edge and re-enter at the opposing side, replicating the scenario where the voxel is surrounded by mirror images of itself on all sides. Given the slow transport of water across a healthy blood-brain barrier over the ∼100 ms timescale simulated here (Herscovitch et al., 1987), vessels were set to be impermeable. Therefore, any MC spin that crossed from IV to EV, or vice versa, would have its new position recalculated until it remained within its original compartment. For our simulation settings, we estimate that less than 0.5% of spins required their positions recalculated per time step, resulting in a negligible increase to the runtime compared to the more computationally intensive Δ*B*_0_ calculations.

For deterministic diffusion, to model impermeable vessels, Eq. [7] was modified to return any magnetization that leaks across boundaries to its original compartment based on convolving the diffusion kernel with either the IV space map (or the EV space map) to derive correction factors for the EV space (or IV space), as described in (Pannetier et al., 2013, 2014). For all approaches, the same diffusion coefficient value was used for the IV and EV spaces.

#### 3.1.4 Signal calculation

Simulations were run with *B*_0_ = 3 T and a single Δ*χ* of 4.15×10^−7^ across all vessels (corresponding to approximately 65% oxygenation and 35% hematocrit, reflecting the venous and capillary weighting of the BOLD signal). Note, the assumption of a single oxygenation level across vessels, while common, is not realistic; however, this assumption was applied across all simulation approaches under matched conditions. A single spin-echo (SE) simulation was run using an echo-time (TE) of 70 ms (refocusing pulse at 35 ms) and a timestep *δt* = 0.2 ms. The gradient-echo (GE) signal was calculated using the same simulations by extracting the signal at TE = 30 ms, before the refocusing pulse.

### 3.2 Simulation approaches

All the simulation approaches that we considered are summarized in Table 2 and the Δ*B*_0_ calculations for the main 2D and 3D geometries are visualized in Fig. 2. Each simulation approach is given a string specifier to indicate the following: the dimensionality of the voxel, the Δ*B*_0_ calculation used, the diffusion calculation used, and any other relevant details. Fig. 3 summarizes the incremental changes between related simulation approaches.

**Fig. 2:**
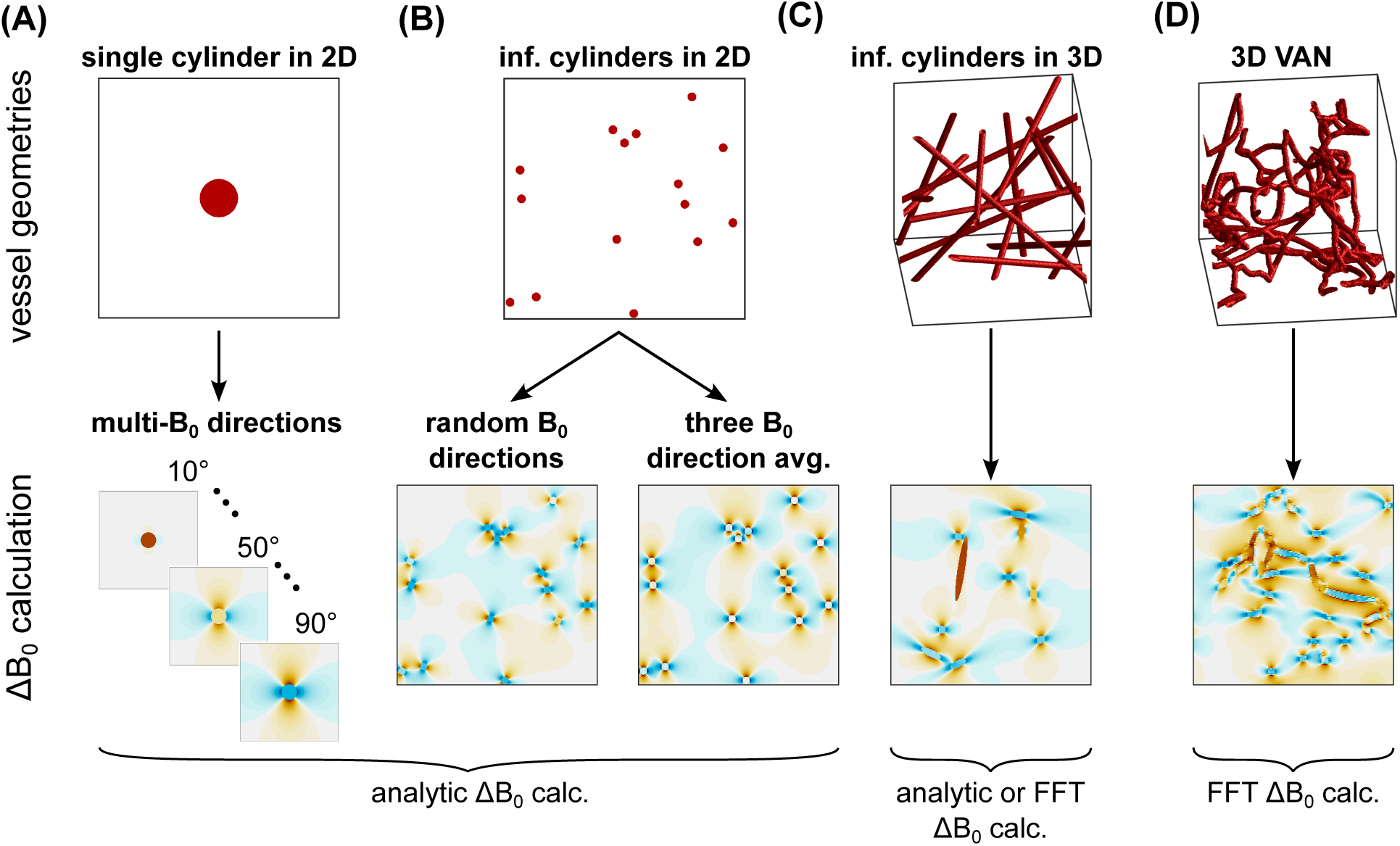
Overview of the vessel geometries and Δ*B*_0_ field offset calculations. The vessel geometries considered were: (A) a single infinite cylinder in 2D where the signal simulations were run at multiple *B*_0_ angles in 10° increments then combined; (B) multiple parallel cylinders in 2D where field offsets were determined based on randomly assigned *B*_0_ directions per vessel or the average of three orthogonal *B*_0_ directions; (C) infinite cylinders randomly distributed and oriented in 3D; and (D) 3D vascular anatomical networks (VANs) composed of tortuous, branching, finite cylinders. The field offset maps show the field offsets across the entire voxel for the 2D geometries or just a representative slice for the 3D geometries. The orange colours indicate increased *B*_0_ field, and the blue colours indicate reduced *B*_0_ field. The labels below the maps indicate whether the simulation comparisons used field offsets calculated analytically or with the FFT-based method. For clarity of display, these vessel networks are smaller than the voxel sizes used for simulation.

**Fig. 3:**
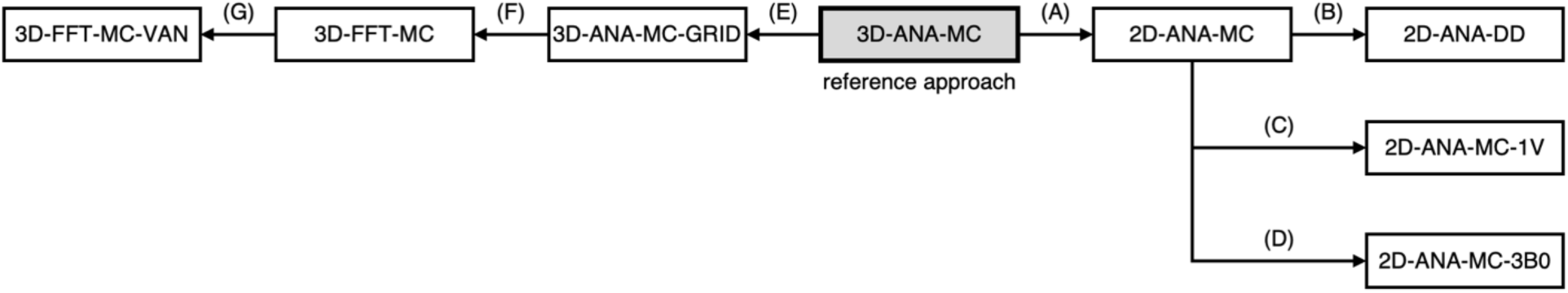
Flow chart depicting the incremental changes between simulation approaches (2D approaches to the right of the reference approach, and 3D to the left). For the 2D methods: (A) 2D-ANA-MC differs from 3D-ANA-MC by tracking spins in a 2D plane where blood vessels are all perpendicular to the simulation plane, and each blood vessel is assigned a randomly oriented *B*_0_ direction. (B) 2D-ANA-DD differs from 2D-ANA-MC by sampling Δ*B*_0_ on a precalculated grid and modelling diffusion using the deterministic diffusion method. (C) 2D-ANA-MC-1V differs from 2D-ANA-MC by using a single vessel perpendicular to the plane. The simulation is repeated over nine equally spaced *B*_0_ angles, and the net signal is given by the weighted sum of signals from all orientations. (D) 2D-ANA-MC-3B0 differs from 2D-ANA-MC by averaging the *B*_0_ offsets from three orthogonal directions instead of randomly assigning the *B*_0_ orientation per vessel. For the 3D methods: (E) 3D-ANA-MC-GRID differs from 3D-ANA-MC by using a precalculated, discretized Δ*B*_0_ map instead of calculating Δ*B*_0_ at each time point for each spin’s position relative to all vessels. (F) 3D-FFT-MC differs from 3D-ANA-MC-GRID by using the Fourier-based Δ*B*_0_ calculation method, not the analytical method. (G) 3D-FFT-MC-VAN differs from 3D-FFT-MC by using finite and branching cylinder segments instead of randomly oriented infinite cylinders.

**Table 2:** Summary of the simulation approaches that were compared. The top half lists the two-dimensional approaches, and the bottom half lists the three-dimensional approaches. DD: deterministic diffusion; FFT: fast Fourier transform; MC: Monte Carlo; VAN: vascular anatomical network

| Name | # dimensions | $\Delta B_0$ calculation | Continuous or Gridded $\Delta B_0$ | Diffusion | Notes |
| --- | --- | --- | --- | --- | --- |
| 2D-ANA-MC <sup>a</sup> | 2 | analytical | continuous | MC | random $B_0$ direction per vessel |
| 2D-ANA-DD <sup>b</sup> | 2 | analytical | gridded | DD | same as above, but using deterministic diffusion |
| 2D-ANA-MC-1V <sup>c</sup> | 2 | analytical | continuous | MC | single vessel, weighted average of simulations over 9 $B_0$ polar angles (10°, 20°, ..., 90°) |
| 2D-ANA-MC-3B0 <sup>d</sup> | 2 | analytical | continuous | MC | average field offset from three $B_0$ directions |
| **3D-ANA-MC <sup>e</sup>** | 3 | analytical | continuous | MC | **reference approach** |
| 3D-ANA-MC-GRID <sub>f</sub> | 3 | analytical | gridded | MC | same as the reference, but $\Delta B_0$ is on a grid |
| 3D-FFT-MC <sup>g</sup> | 3 | FFT | gridded | MC | same as above, but $\Delta B_0$ calculated using FFT |
| 3D-FFT-MC-VAN <sup>h</sup> | 3 | FFT | gridded | MC | simplified VANs using finite branching vessels (single diameter) with realistic tortuosity |

Miller & Jezzard (2008). <sup>b</sup>Berman et al. (2018). <sup>c</sup>Ogawa et al. (1993) and Uludag et al. (2009). <sup>d</sup>Pannetier et al. (2013). <sup>e</sup>Boxerman, Hamberg, et al. (1995). <sup>f</sup>X. Cheng et al. (2019). <sup>g</sup>Pathak et al. (2008). <sup>h</sup>Marques & Bowtell (2008), Pathak et al. (2008), and Gagnon et al. (2015).

#### 3D-ANA-MC: reference simulation approach

The reference simulation approach that all other simulations were compared against was based on the seminal work of Boxerman, Hamberg, et al. (1995). These simulations consisted of a 3D voxel with randomly-oriented and randomly-positioned infinite cylinders using analytically calculated *B*_0_ offsets and MC-based diffusion in continuous space. The field offset experienced by each spin at each timestep was calculated for each spin’s position relative to each vessel. This combination of simulation options was chosen for the reference approach since it is commonly implemented and replicated in studies and may be considered the most accurate in terms of the approximation of the diffusion process (as opposed to deterministic diffusion), and the accuracy of the *B*_0_ calculation does not suffer from the Fourier-based Δ*B*_0_ gridding artifacts. When comparing the reference to VAN simulations, the results are not necessarily expected to match due to morphological differences likes vessel branching, tortuosity, and asymmetries in the synthetic VAN capillary networks; therefore, differences between the VAN simulations and the reference should not necessarily be classified as “errors”.

Ten voxels were used, each with a unique set of vessels. All other simulation approaches that used infinite cylinders in 3D (i.e., 3D-ANA-MC-GRID and 3D-FFT-MC) used the same ten voxels. The orientation of each vessel was determined by the polar and azimuthal angles of the cylinders. The polar angle of each cylinder was selected from a sin(*θ*) distribution, which was implemented by choosing *θ* = acos(2*u* – 1), where *u* was randomly drawn from the uniform distribution [0,1). The azimuthal angle was randomly drawn from the uniform distribution [0, 2π). To spatially distribute cylinders uniformly throughout the voxel, once the cylinder direction was determined, a point that the cylinder axis passes through was randomly selected on the plane orthogonal to the cylinder direction and passing through the origin of the voxel (Martindale et al., 2008). Cylinders were added in this way until the desired blood volume fraction was reached.

#### 2D-ANA-MC

In 2D, the blood volume fraction can be easily controlled if all vessels are perpendicular to the simulation plane. Unlike in 3D, where vessels are randomly oriented and the direction of *B*_0_ is fixed, one way of generating a realistic distribution of field offsets in 2D is to use a fixed orientation of vessels (perpendicular to the plane) but assign a random *B*_0_ direction to each vessel (Berman et al., 2018; Miller & Jezzard, 2008). The *B*_0_ orientation per vessel was determined using the same distributions for the polar and azimuthal angles as described for the vessel orientation in the 3D-ANA-MC method above. The 2D-ANA-MC simulations randomly and uniformly positioned vessels in the plane (appearing as circles, as in Fig. 2B), distributed their *B*_0_ directions as mentioned, and used analytical *B*_0_ calculations and MC diffusion (Miller & Jezzard, 2008).

#### 2D-ANA-DD

We repeated simulations using a similar configuration as 2D-ANA-MC but with deterministic diffusion (Berman et al., 2018). By necessity, the *B*_0_ offsets and magnetization were calculated on a grid. The diffusion kernel spanned from –6*σ* to +6*σ*. To ensure that the kernel did not behave like a delta function, the spatial resolution of the grid was chosen such that the diffusion kernel contained a minimum of 13 elements. Since the effective voxel size was scaled to model different diameters, the largest vessel radius, 60-μm, imposed the strictest constraint on the kernel. A 15,860×15,860 grid size satisfied the kernel size requirement in this case. For smaller radii, *R*, the number of grid elements per side was linearly scaled as 15,860×*R* [μm]/(60 μm) and rounded up to the nearest multiple of ten. A minimum of 1000 elements per side was imposed since going below this value resulted in an inadequate sampling of the vessels and their field offsets, which was qualitatively apparent when comparing simulations across resolutions (results not shown). Therefore, simulations with radii < 4 μm used 1000 elements per side. To account for convolution contributions from outside the voxel, the magnetization of the voxel was padded using a wrapped version of itself, similar to how spins wrap at the boundaries during MC diffusion.

#### 2D-ANA-MC-1V

These simulations were in 2D using analytically calculated field offsets and MC diffusion. Following seminal modelling studies, a single vessel was placed at the centre of a 2D voxel and oriented perpendicular to the plane (Ogawa et al., 1993; Uludag et al., 2009). Simulations were repeated with the *B*_0_ polar angle increased in 10° increments from 10° to 90° and the net signal was calculated as the weighted average over the simulation from each angle (*S_θ_*):

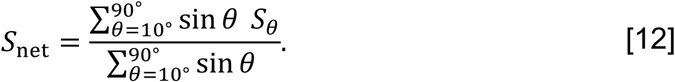

Since there was no randomness associated with the vasculature, simulations were performed on a single voxel, resulting in no estimated uncertainty. We use “1V” in the label to denote a single vessel.

#### 2D-ANA-MC-3B0

These simulations were similar to 2D-ANA-MC, however, the *B*_0_ offsets for the vessels were calculated using the average of three orthogonal *B*_0_ directions: one parallel and two perpendicular to the vessels (Pannetier et al., 2013). In practice, since the EV offsets of the two perpendicular orientations may cancel each other out if they are perpendicular to each other, a single perpendicular orientation was used, and the net field offset was the weighted sum of 1/3 the parallel and 2/3 the perpendicular offsets. For consistency with the original description of the technique, we continue to refer to this as the “3*B*_0_” approach. The *MrVox2D* simulation toolkit uses the 3*B*_0_ approach, although with Fourier-based field calculations and deterministic diffusion (Pannetier et al., 2013).

#### 3D-ANA-MC-GRID

Fourier-based simulation approaches perform calculations on a discretized grid, making comparison with the reference approach potentially ambiguous regarding whether the source of any differences originates from the calculation of Δ*B*_0_ using the Fourier transform or from the discretization of space. Therefore, to bridge between the reference simulations (3D-ANA-MC) and the Fourier-based approaches, simulations were performed that were similar to 3D-ANA-MC, with the exception that Δ*B*_0_ was calculated on a regularly spaced grid by summing the analytical Δ*B*_0_ contributions from every vessel using Eq. [1] at every grid point (X. Cheng et al., 2019). Therefore, rather than updating the field offset that each spin experiences relative to every vessel, the spin position is used to retrieve the precomputed field offset from the discretized grid. Note, for this and the remaining MC simulations, the spin positions were computed in continuous space throughout the voxel, despite Δ*B*_0_ being computed in discrete space.

#### 3D-FFT-MC

This simulation applied the Fourier-based field offset calculation to infinite cylinders on the same grid as in the previous method (i.e., 3D-ANA-MC-GRID) using Eq. [4] (Pathak et al., 2008). The susceptibility distribution was discretized by analytically creating a vessel mask based on whether the centre of each grid point was in a vessel or not. To model the effect of surrounding vasculature, cyclic convolution was calculated in *k* space with the dipole kernel size equal to the grid size (i.e., the voxels were not padded before the application of the Fourier transform). We label this and the following approach with “FFT” since the fast Fourier transform algorithm was used to calculate the Fourier transforms.

#### 3D-FFT-MC-VAN

The final simulation approach was based on vascular anatomical networks (VANs) (Gagnon et al., 2015; Marques & Bowtell, 2008; Pathak et al., 2008). We synthesized a 1-mm^3^ isotropic VAN that consisted of a capillary bed of tortuous vessels made from smoothly connected, finite cylinders without overlap. To enable fair comparison with the infinite cylinder model, a single vessel radius (2.85 µm) was assigned to all vessels in the VAN. Due to the fixed radius, and to achieve equivalent CBV as the infinite cylinder models, we increased the number of capillary segments to 26,100 segments/ml (compared to an average of 11,470 segments/ml from reconstructed VANs of equivalent size (Hartung, Badr, Mihelic, et al., 2021)—a factor of 2.25× higher vessel density). These VANs were constructed with otherwise realistic vascular synthesis methods that offer realistic branching patterns and individual segment distributions of length and tortuosity (Hartung, Badr, Moeini, et al., 2021; Hartung et al., 2025; Linninger et al., 2019). Pial vessels and perpendicular diving/draining vessels, which are typically much larger, were removed from the VAN after synthesis. We refer to these networks as *simplified* VANs, due to the removal of the larger vessels and forcing all capillaries to a single radius. To produce the 10 voxels to run simulations on, a large simplified VAN was created using the described methods, and then it was divided into ten smaller VANs of 400-μm voxel edge length. Vessels of different radii, from 1 to 60 μm, were simulated by changing the apparent voxel size, as described in the *Voxel/geometry definition* section above (giving an effective voxel edge length of 140 μm when the radius was 1 μm). Δ*B*_0_ was calculated on these smaller voxels using the Fourier-based method with cyclic convolution. For full VANs, we recommend non-cyclic convolution along the direction perpendicular to the pial surface to ensure that field offsets from the large pial vessels do not wrap around to the lower depths of the voxel. More details on how the vessel masks were generated for the Δ*B*_0_ calculations are provided in the Supplementary Material. The resulting blood volume of the synthesized VANs was slightly below the nominal volume of the infinite cylinder networks (1.79% vs. 2%); therefore, to compensate, we applied a scaling factor to the simulated VAN signals, as proposed by Kiselev & Posse (1999). Further details on this volume mismatch and the signal scaling are detailed in the Supplementary Material.

### 3.3 Analyses

The analysis was structured to address the guiding questions from the Introduction. All simulations were first averaged across all ten voxels, and then their relative accuracy was compared by three different metrics. First, as a relevant application where BOLD simulations may be applied, we performed a simplified vascular fingerprinting analysis to determine the impact of the simulation approach on the estimated vessel radius. This was followed by in-depth simulation accuracy analyses. The simulated signals remained complex-valued quantities during the analyses, except where specified. Finally, the computational performance of all simulations was evaluated.

#### 3.3.1 Use case: vascular fingerprinting accuracy

MR vascular fingerprinting is a technique that analyzes the time evolution of the MR signal to estimate quantitative information about the microvasculature, such as mean vessel radius and blood oxygenation, based on matching the measured signal to a lookup table of precalculated simulated signals (Christen et al., 2014). In our simplified vascular fingerprinting analysis, each simulation approach was used to generate a signal dictionary from which the vessel radius was to be extracted from the “measured” test signal (3D-ANA-MC, i.e., the reference approach). The dictionary and test signals used the evolution of the simulated signals as a function of time at all simulated time points. The total (EV+IV) signals were normalized by their temporal average, then the test signals were compared to the dictionary signals across all 15 radii (1–60 μm) using the coefficient of determination, *R*^2^ (Christen et al., 2014). The radius that produced the dictionary signal with the highest *R*^2^ was selected as the estimated radius. This analysis was repeated with the reference simulations from all 15 radii used as the test signals. While this simplified fingerprinting analysis is not implemented exactly as experimental fingerprinting experiments have been (e.g., limited range of fit parameters, test signals drawn from noise-free simulations rather than experimentally measured, and voxel sizes that depend on vessel radius), this analysis is specifically meant to identify biases that may be introduced by the choice of simulation technique per se when applied to quantitative physiological imaging, as per Guiding Question #1.

#### 3.3.2 Simulation accuracy: comparison of signal intensity

To understand the outcome of the fingerprinting analysis, we further compared specific attributes of the signal evolutions pertaining directly to simulation accuracy, starting with signal intensity. The average signal evolution for each simulation approach was compared to the average reference simulations (3D-ANA-MC) using the normalized root mean square deviation (NRMSD), defined as

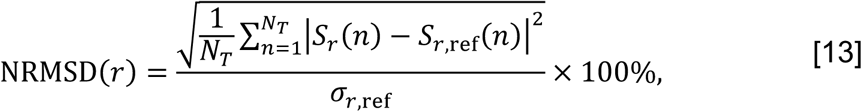

where *r* indexes radius, *N_T_* is the number of time points in the simulations, *S_r_* is the mean simulated signal evolution for the *r*-th radius, *S_r_*_,ref_ is the mean signal evolution from the reference method for the *r*-th radius, and *σ_r_*_,ref_ is the standard deviation over time of the reference signal evolution for the *r*-th radius. NRMSD was calculated separately for the total (EV+IV), EV, and IV signals.

#### 3.3.3 Simulation accuracy: comparison of BOLD contrast

Next, the apparent GE and SE relaxation rates were calculated at a single TE as

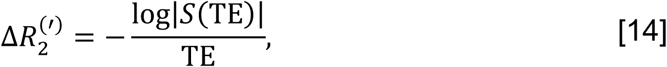

using TE = 30 ms for Δ*R*_2_*’* and TE = 70 ms (i.e., the spin-echo time) for Δ*R*_2_. Plots of the relaxation rates vs. vessel radius were used to characterize the simulations. We refer to these plots as “Boxerman plots” given their use in the seminal work of Boxerman, Hamberg, et al. (1995).

A range of factors, such as the field offsets themselves (in the case of static dephasing) or the correlation of the field offsets sensed by the spins over time (in the case of motional narrowing) can influence how the relaxation rates may differ from the reference simulation. Therefore, to aid in the interpretation of the results, the distributions of *B*_0_ offsets were analyzed for a subset of the 2D and 3D geometries. Field offsets across each voxel were calculated on a grid (1000^2^ in 2D and 500^3^ in 3D) from which histogram distributions were generated.

#### 3.3.4 Computational Performance

Finally, we previously reported computational performance of most—but not all—of the simulation approaches assessed here (Chaussé et al., 2025). To assess the remaining approaches, sample simulations were run from which the peak memory demands and the total computing time were recorded. The computing time was divided into the *initialization time* for the simulations (e.g., create the set of vessels or, if necessary, calculate the Δ*B*_0_ maps) and the *run time* to calculate the MR signal. More details can be found in the Supplementary Material.

## 4 Results

### 4.1.1 Use case: vascular fingerprinting accuracy

Summarized in Fig. 4 are the comparisons of the outcomes of our vascular fingerprinting use case. We focus specifically on the absolute difference (i.e., absolute error) between the estimated and true vessel radius for all simulation approaches. Box plots of the resulting absolute error in the estimated radii are shown in Fig. 5. Except for 2D-ANA-MC-3B0, which resulted in errors in 12 of the 15 radii (mean/max absolute errors of 54%/233%), and 2D-ANA-MC-1V, which produced errors at seven radii (mean/max absolute errors of 13%/33%), most approaches could reasonably predict the ground-truth radii (see Fig. 5). The Fourier-based approaches, including the 3D-FFT-MC and 3D-FFT-MC-VAN simulations, only had two erroneous radii each. The remaining four approaches predicted all radii correctly.

**Fig. 4:**
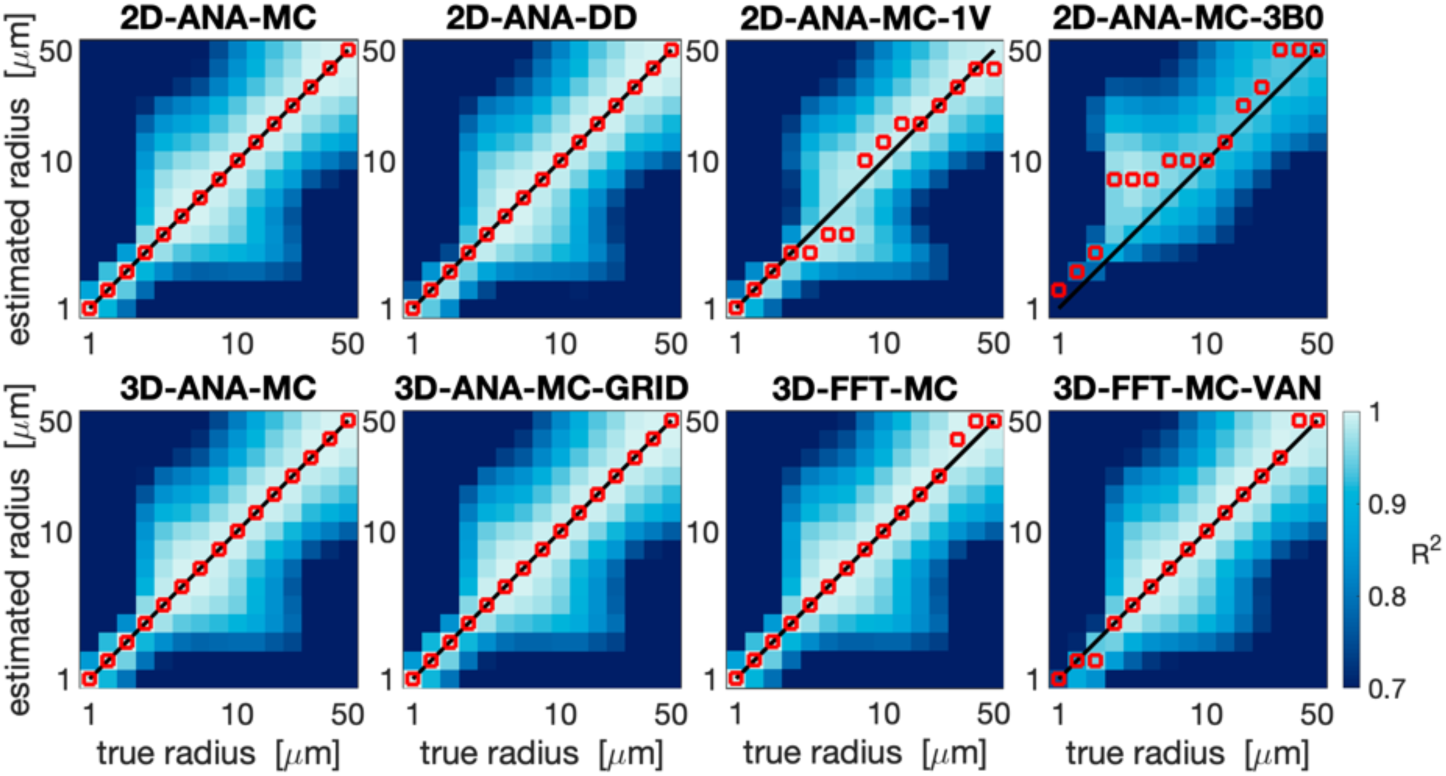
Vascular fingerprinting results showing the coefficient of determination, *R*^2^, when each respective simulation approach is used as the fingerprinting dictionary. The horizontal axes show the true radius corresponding to the test signals from the 3D-ANA-MC simulations, and the vertical axes show the dictionary radius used for comparison. The solid black line shows the ground-truth one-to-one relationship, and the red markers show the estimated radius corresponding to the dictionary radius with the highest *R*^2^ for the given test signal. The plots are organized by the 2D approaches in the top row and the 3D approaches in the bottom row. All plots share the colour bar in the bottom right.

**Fig. 5:**
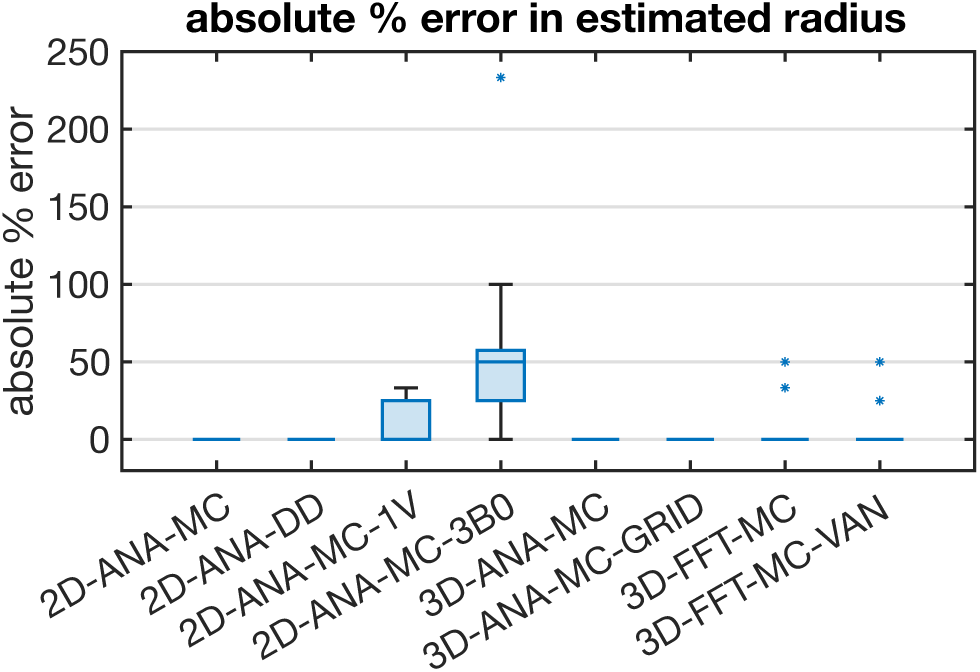
Box plot of the absolute percent error in the radius estimated using the dictionary lookup relative to the true radius from the 3D-ANA-MC simulations. The median absolute error is represented by the central line, the interquartile range is represented by the box height, the minimum and maximum non-outlier values are represented by the whiskers, and any outliers are represented by the * symbols. Several simulation approaches gave perfect estimates for all radii and, hence, are represented by a single horizontal line at zero error.

### 4.1.2 Simulation accuracy: comparison of signal intensity

To interpret the outcomes of the use case test, the most fundamental comparison was that of the signal intensities. Differences in intensity were computed relative to the reference method (i.e. 3D-ANA-MC) to produce the NRMSD. The NRMSD of all simulation techniques are plotted as a function of radius in Fig. 6, and the average NRMSD across radii is shown in Fig. 7. The associated mean simulated SE signal evolution for each simulation technique is shown at three characteristic radii in the Supplementary Material Fig. S2. The EV signal for all approaches agreed closely with the reference for most radii, although many techniques still had poorer agreement at smaller radii (Fig. 6). The IV signal was typically associated with the greatest relative difference (∼10% or greater) across all techniques. While a 10% difference may seem high, from the example IV signal evolutions in Supplementary Material Fig. S2, the approaches with NRMSD ≲ 10% are still in excellent agreement overall with the reference signal (e.g., 2D-ANA-MC, 3D-ANA-MC-GRID). Given the low blood volume (2%), the total signal and its NRMSD were dominated by the EV signal. An exception to this is the 2D-ANA-MC-3B0 simulations, where the total signal was extremely biased due to the erroneous IV signal. Of note, the EV and total signal from the VAN simulations in Fig. 6G have NRMSD < 10% for radii ≳ 3 μm, in line with gridded infinite cylinders (3D-ANA-MC-GRID), giving acceptable results when qualitatively assessed, as shown in the Supplementary Material Fig. S2G, where the signal evolutions overlap closely for the 10-μm and 60-μm radius simulations.

**Fig. 6:**
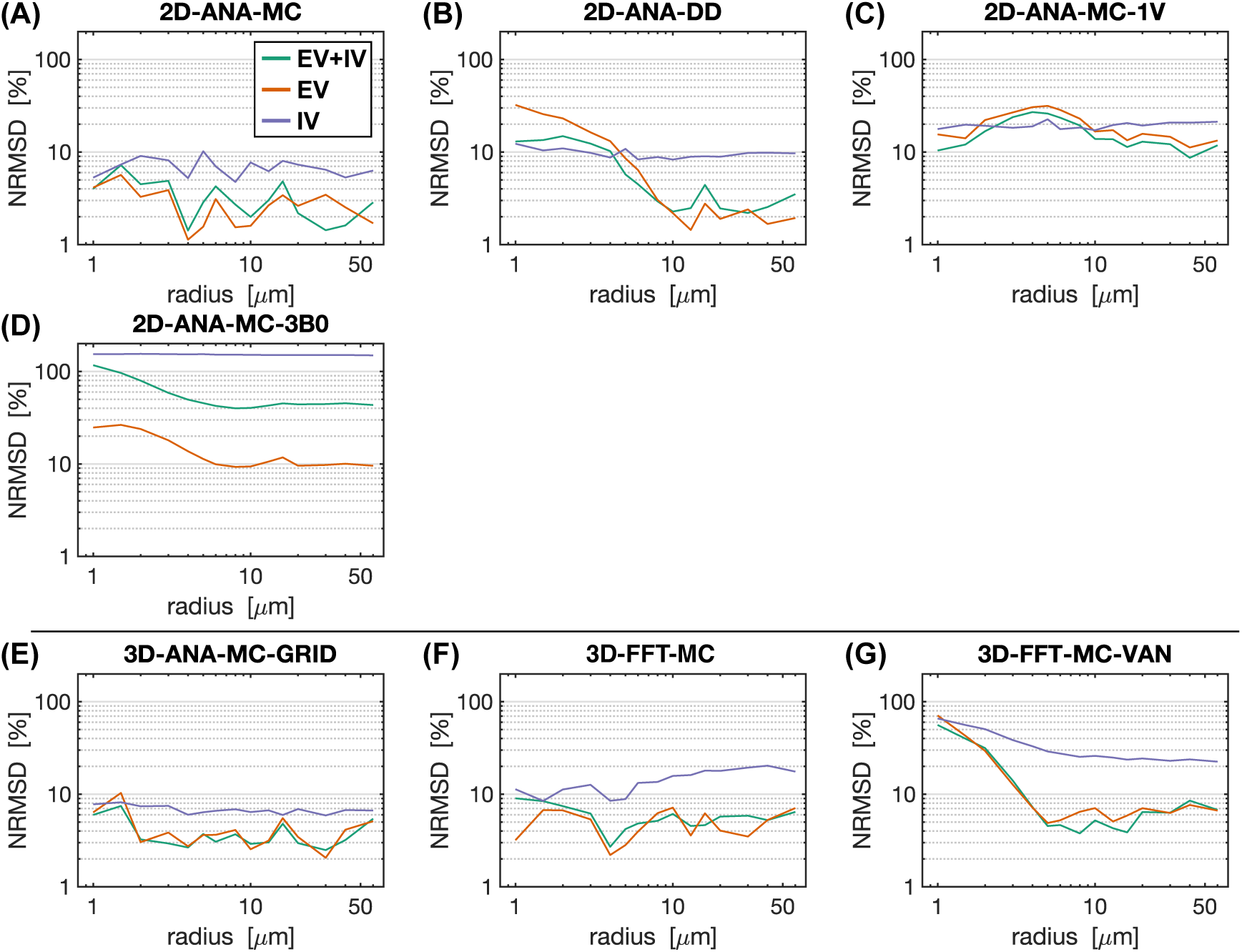
Normalized RMS difference (NRMSD) between the mean simulated decay for each simulation technique relative to the mean reference simulation (3D-ANA-MC) vs. radius. (A)–(D) NRMSD of the 2D voxel simulations. (E)–(F) NRMSD of the 3D voxel simulations. NRMSD was calculated separately for the total signal (EV+IV, green curves), EV only (orange), and IV only (purple). Note that all graphs are plotted on a log-log scale.

**Fig. 7:**
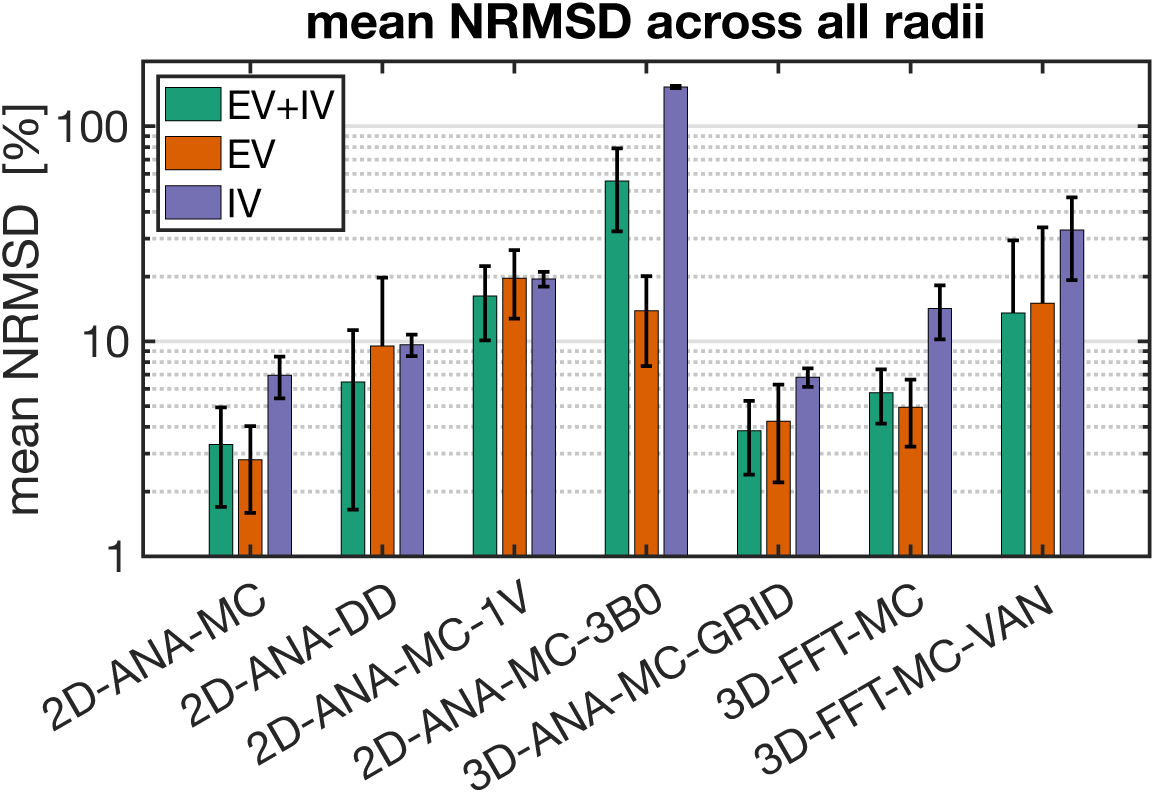
Bar plots of the normalized RMS difference (NRMSD) averaged over all radii for each simulation technique relative to the reference technique (3D-ANA-MC). The error bars represent ± one standard deviation of the mean. In cases where the standard deviation is greater than or equal to the mean, only the upper bar is displayed since values are displayed on a logarithmic scale. NRMSD is plotted for the total signal (EV+IV, green bars), EV only (orange), and IV only (purple).

As an attempt to remove biases that may be imposed by selecting the 3D analytical infinite cylinder MC approach as the reference, we repeated the analyses in Fig. 7 using each of the simulation approaches for the reference signals. Those results are shown in Supplementary Material Fig. S3. From those plots, the trends that were observed when 3D-ANA-MC was the reference signal generally persisted when the other approaches were used as reference signals. The 2D-FFT-MC-3B0 approach is a clear outlier when it was used as the test or reference signals. When 3D-FFT-MC-VAN was used for the reference signals (bottom row), the NRMSD values were slightly elevated, particularly for the IV signal, but still in line with the NRMSD of other approaches, such as 2D-ANA-DD.

### 4.1.3 Simulation accuracy: comparison of BOLD contrast

The further quantify the signal evolutions, we compared the simulations in terms of the vascular contributions to the *R*_2_ (SE) and *R*_2_*’* (GE) relaxation rates. The Boxerman-style plots comparing the relaxation rates as a function of radius are shown in Fig. 8 for all 2D methods. Different diffusion techniques (Monte Carlo and deterministic diffusion) are compared in Fig. 8A. The 2D-ANA-MC and 2D-ANA-DD techniques both showed excellent agreement with the 3D-ANA-MC reference approach, although at small radii, the IV SE relaxation rates for both approaches diverged in opposite directions (Fig. 8A.4). Simplifying the simulations using a single vessel is shown in Fig. 8B. The 2D-ANA-MC-1V approach slightly overestimated the EV relaxation rates (Fig. 8B.2), underestimated the IV GE rate (Fig. 8B.3), and was unable to predict the IV SE rate (Fig. 8B.4) since the IV space is completely homogeneous; hence, spins are refocused at every *B*_0_ angle. The use of a random *B*_0_ direction per vessel (2D-ANA-MC) is compared to three *B*_0_-direction averaging (2D-ANA-MC-3B0) in Fig. 8C. The 2D-ANA-MC-3B0 technique slightly underestimated the EV relaxation rates (Fig. 8C.2). The IV GE relaxation rates (Fig. 8C.3) were all nearly zero since all vessels were defined as having the same IV Δ*B*_0_ value in the 3B0 approach, although there is still a small influence from the neighbouring EV fields on the IV space, resulting in a dramatic reduction in the total relaxation rate. Yet, the IV SE rates for the 3*B*_0_ approach (Fig. 8C.4) are consistent with the 2D-ANA-MC method since Δ*R*_2_ is dominated by EV fields.

**Fig. 8:**
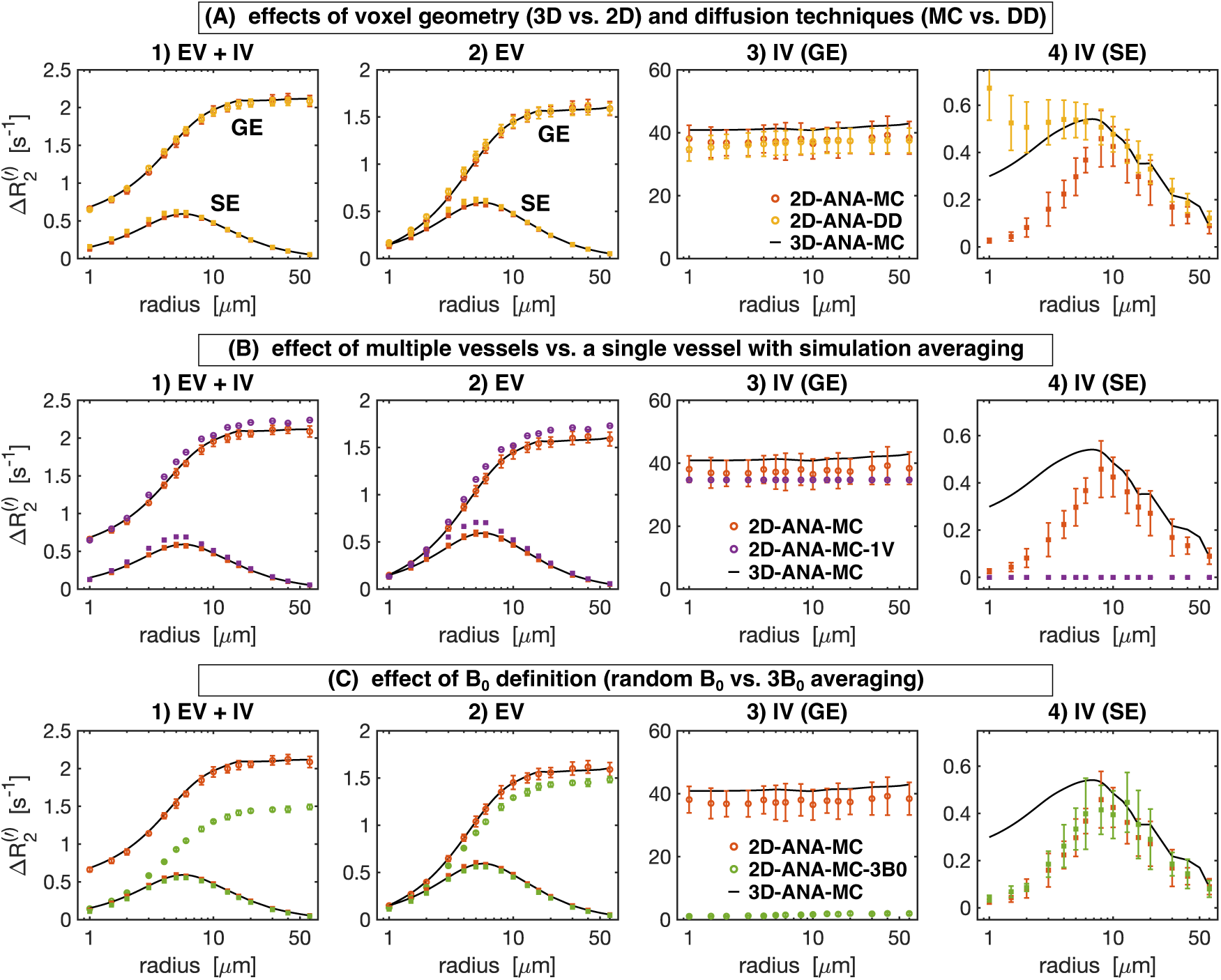
Boxerman plots showing the gradient-echo (GE) and spin-echo (SE) relaxation rates (Δ*R*_2_^(^*’*^)^) as a function of vessel radius for the 2D geometry simulation techniques. From left to right: relaxation rates are plotted separately for 1) total signal (EV+IV), 2) extravascular (EV), 3) intravascular GE, and 4) intravascular SE. In all plots, the reference relaxation rates from 3D-ANA-MC are displayed as black curves after applying a smoothing spline fit to them, and the techniques being compared use circle symbols for Δ*R*_2_*’* and square symbols for Δ*R*_2_. (A) Comparison between simulations modelling diffusion using the Monte Carlo method (2D-ANA-MC) or deterministic diffusion (2D-ANA-DD). The 2D-ANA-MC technique is expected to be closest approximation to the reference 3D-ANA-MC technique (black lines), hence the comparison of 3D to the simplified 2D models. (B) Comparison between simulations from voxels filled with multiple vessels (2D-ANA-MC) to the average simulations from a voxel occupied by a single vessel and uniformly varying *B*_0_ polar angle. (C) Comparison between simulations using field offsets generated by random *B*_0_ directions per vessel (2D-ANA-MC) to the average offset from three *B*_0_ directions per vessel (2D-ANA-MC-3B0). Error bars show the mean ± standard deviation across all simulations.

The Boxerman-style plots for all 3D methods are shown in Fig. 9. The effect of analytical vs. Fourier-based Δ*B*_0_ calculations is compared in Fig. 9A. Except for the case of IV SE (Fig. 9A.4), the Δ*R*_2_^(^*’*^)^ curves for both Δ*B*_0_ calculation approaches are in excellent agreement with that of the reference technique. Discretization alone (comparing 3D-ANA-MC-GRID with 3D-ANA-MC) caused the IV SE relaxation rates (Fig. 9A.4) to increase at small radii relative to the reference. Conversely, when Δ*B*_0_ is calculated on a grid using the Fourier method (3D-FFT-MC), the simulated relaxation rates associated with the larger radii are the least accurate, and the entire Boxerman curve appears to shift toward larger radii with higher relaxation rates. Fig. 9B compares the effect of simulating with infinite cylinders to VANs, both using Fourier-based Δ*B*_0_ calculations. The total and EV signal relaxation rates are in good agreement with each other and the reference (Fig. 9B.1 and Fig. 9B.2), and the IV relaxation rates are also in good agreement until the radius falls below ∼20 μm, at which point the IV relaxation rates for the VANs start to depart from both infinite cylinder cases, i.e., 3D-ANA-MC and 3D-FFT-MC (Fig. 9B.3 and Fig. 9B.4).

**Fig. 9:**
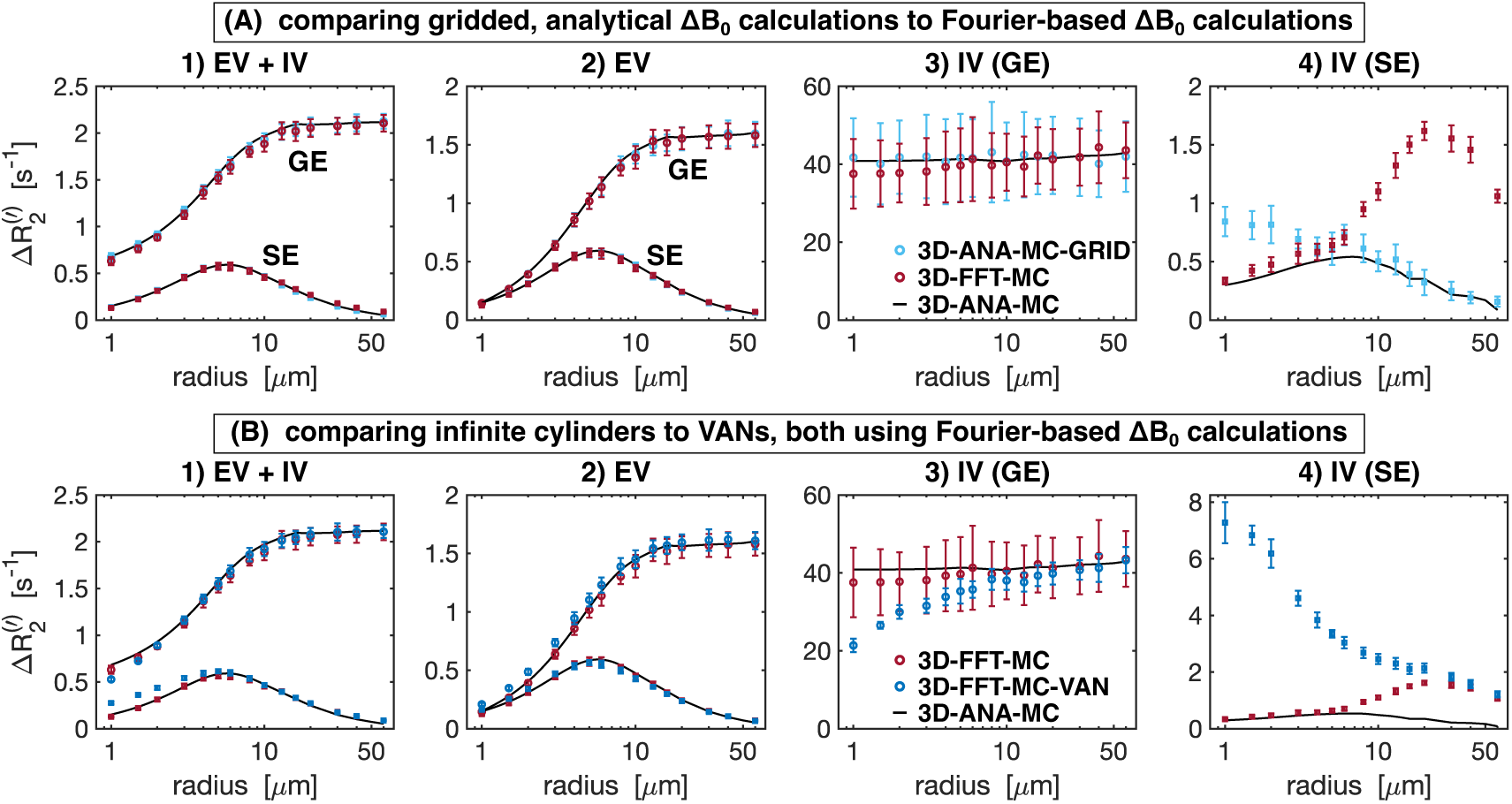
Boxerman plots showing the gradient-echo (GE) and spin-echo (SE) relaxation rates (Δ*R*_2_^(^*’*^)^) as a function of vessel radius for the 3D geometry simulation techniques. From left to right: relaxation rates are plotted separately for 1) total signal (EV+IV), 2) extravascular (EV), 3) intravascular GE, and 4) intravascular SE. In all plots, the reference relaxation rates from 3D-ANA-MC are displayed in black after applying a smoothing spline fit to them, and the techniques being compared use circle symbols for Δ*R*_2_*’* and square symbols for Δ*R*_2_. (A) The effect of gridding the analytical field offsets (3D-ANA-MC-GRID) is compared to calculating the offsets using the Fourier method on infinite cylinders (3D-FFT-MC). (B) The VAN simulations are compared against infinite cylinder simulations where each used the Fourier method. Error bars show the mean ± standard deviation across all simulations.

To aid in the interpretation of the preceding differences in the relaxation rates, histograms of the Δ*B*_0_ distributions for selected 2D and 3D approaches are shown in Fig. 10 and Fig. 11, respectively. The more dissimilar the Δ*B*_0_ spectra of any of the approaches are, the less likely it is for the simulated signals to match. Conversely, even if Δ*B*_0_ spectra match, the simulated signals may not necessarily also match since the spatial pattern of field offsets impacts the final signal due to the diffusion of water through the EV and IV field offsets. For the 2D geometries, the total and EV Δ*B*_0_ distributions for randomly assigned *B*_0_ directions and the 3*B*_0_ technique are in close agreement with the distributions from 3D infinite cylinders (Fig. 10A, B). The IV histograms from the 3D and 2D (with random *B*_0_ directions) approaches are broad distributions coming to a sharp peak at −67 ppb (Fig. 10C). However, with the 3*B*_0_ technique, since all vessels effectively only see the sum of two *B*_0_ directions, there is an artifactually narrow peak in its IV Δ*B*_0_ distribution. For the 3D techniques, we considered histograms for infinite cylinders with Δ*B*_0_ calculated analytically or with the FFT and VANs using the FFT (Fig. 11). The EV distributions are all remarkably similar (Fig. 11B). When calculated with the FFT, the IV histogram of the infinite cylinders becomes a blurred version of the analytical histogram with a reduced peak at −67 ppb (Fig. 11C), and the IV VAN FFT histogram closely resembles it. Despite the IV spectra for the VAN and infinite cylinder FFT approach being so similar, the simulated signals are quite different (e.g., Fig. 9B.4 and Supplementary Material Fig. S2), confirming that differences in the spatial distributions of Δ*B*_0_, due to tortuosity, vessel junctions, etc., must also be influencing the signals.

**Fig. 10:**
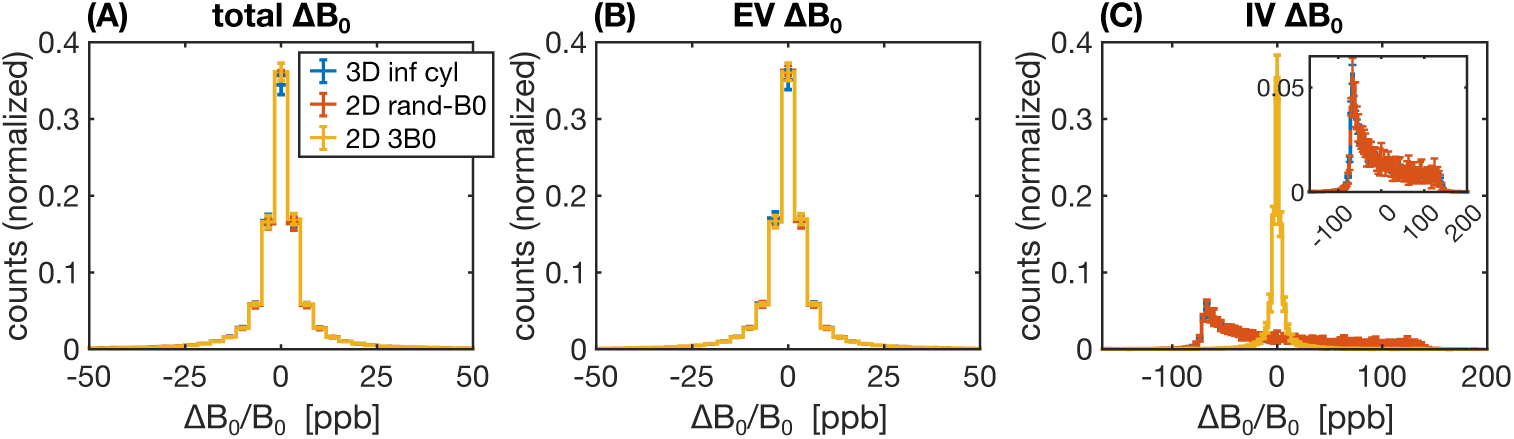
Histograms of the relative Δ*B*_0_ distributions (in parts per billion, ppb) for 2D geometries relative to the reference 3D geometry, considering Δ*B*_0_ over (A) all space, (B) the EV space, or (C) the IV space. The three histograms considered were for random infinite cylinders in 3D (blue), infinite cylinders in 2D with the direction of *B*_0_ randomly assigned per vessel (orange), and infinite cylinders in 2D using the 3*B*_0_ technique (yellow). The inset in (C) shows a zoom-in of the IV histograms, excluding the 3*B*_0_ technique. In all cases, field offsets were calculated analytically, and each histogram is the average of ten voxels’ histograms, with the error bars representing plus/minus one standard deviation. (A)–(C) share the legend in (A).

**Fig. 11:**
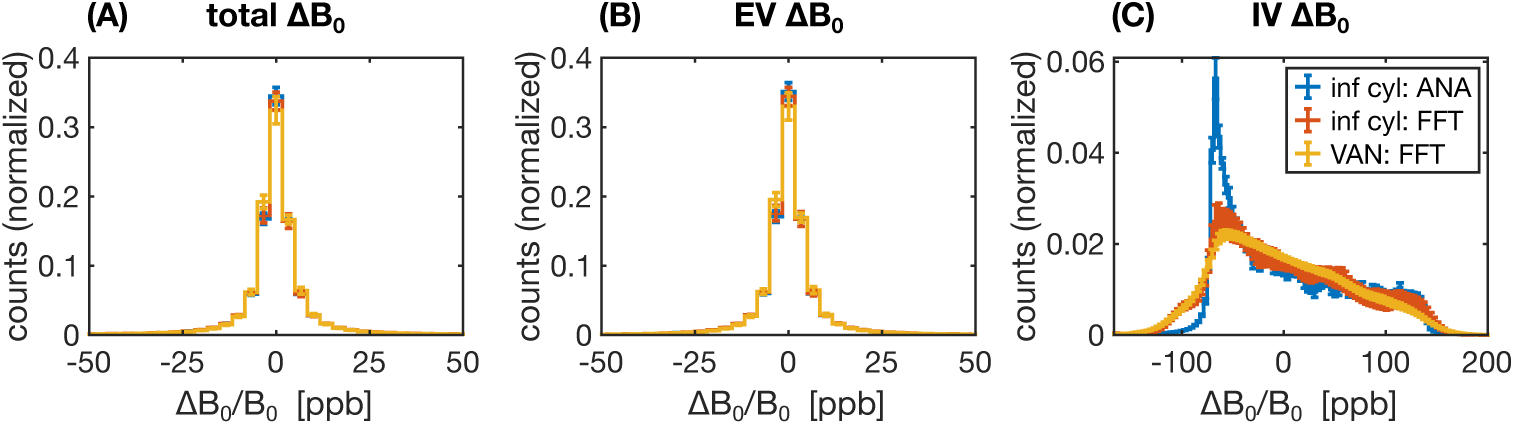
Histograms of the relative Δ*B*_0_ distributions (in parts per billion) for three different combinations of 3D vessel type and *B*_0_ calculation, considering Δ*B*_0_ over (A) all space, (B) the EV space, or (C) the IV space. The three combinations considered were random infinite cylinders with ΔB_0_ calculated analytically (blue) or calculated using the FFT method (orange) or VANs using the FFT method (yellow). In all cases, each histogram is the average of ten voxels’ histograms, with the error bars representing plus/minus one standard deviation. (A)–(C) share the legend in (C).

### 4.1.4 Computational performance

Finally, as mentioned earlier, a key factor in choosing the simulation approach has been computational complexity. The peak memory demands and the total computing time of the simulations are given in Supplementary Material Fig. S4. The reference approach (3D-ANA-MC) had low memory requirements and relatively short computing times. The 2D approaches naturally had the shortest computing times, and the gridded 3D approaches had the longest computing times and greatest memory requirements, as one might expect. One exception to this, however, was for 2D-ANA-DD, where the computational requirements scaled with vessel size due to the necessity to maintain a higher effective spatial resolution that could adequately sample the diffusion kernel. The resulting computing times and memory requirements for 2D-ANA-DD at the largest radii were comparable to the 3D gridded approaches (see Fig. 9 in Chaussé et al. (2025) for more details). Therefore, if simulating on a standard desktop computer with <32 GB of memory, selecting a 2D or 3D analytical Δ*B*_0_ method without gridding should perform well. Overall, the reference approach provided an excellent balance between computational efficiency and numerical accuracy.

## 5 Discussion

Our study has provided a comprehensive comparison of the most common biophysical simulation techniques for modelling the BOLD fMRI signal, including different geometries and implementation choices. A simplified vascular fingerprinting experiment was used to demonstrate how the choice of simulation approach may impact a study’s conclusions (vessel radius, in this case). To provide insight into this impact, we showed that most of the simulation techniques were in good agreement with the reference technique (3D-ANA-MC) in terms of NRMSD and relaxation rates; however, some notable exceptions were found for a subset of the approaches, particularly for the IV component. These findings can provide a baseline for understanding the effect of the choice of simulation approach on the accuracy of quantitative fMRI techniques.

### 5.1 Comparing simulation approaches

With our Guiding Questions in mind, we performed the following analyses:

1. To determine how relevant, modern applications may be impacted by the choice of simulation method, we examined the estimated vessel radii in a simplified vascular fingerprinting analysis. A small subset of approaches was less appropriate for this chosen application.
2. To determine how different infinite cylinder simulation approaches compare to each other, we compared them all to a common reference approach, which we deemed the ground truth for infinite cylinder simulations. See below for more specific comparisons.
3. To determine the effect of assuming blood vessels are infinite cylinders, we directly compared the simulations from synthetic VANs with our reference infinite cylinder model (3D-ANA-MC). To probe the sources of the differences between the reference and VAN simulations, we incrementally changed the simulation approaches from continuous space to discrete space, then from analytical to Fourier-based Δ*B*_0_ calculations, and then from infinite cylinders to VANs. All these simulations were in reasonable agreement (see below for further discussion).

We summarize the impact of the major design decisions when selecting a simulation approach in Table 3 and in the discussion that follows.

**Table 3:**
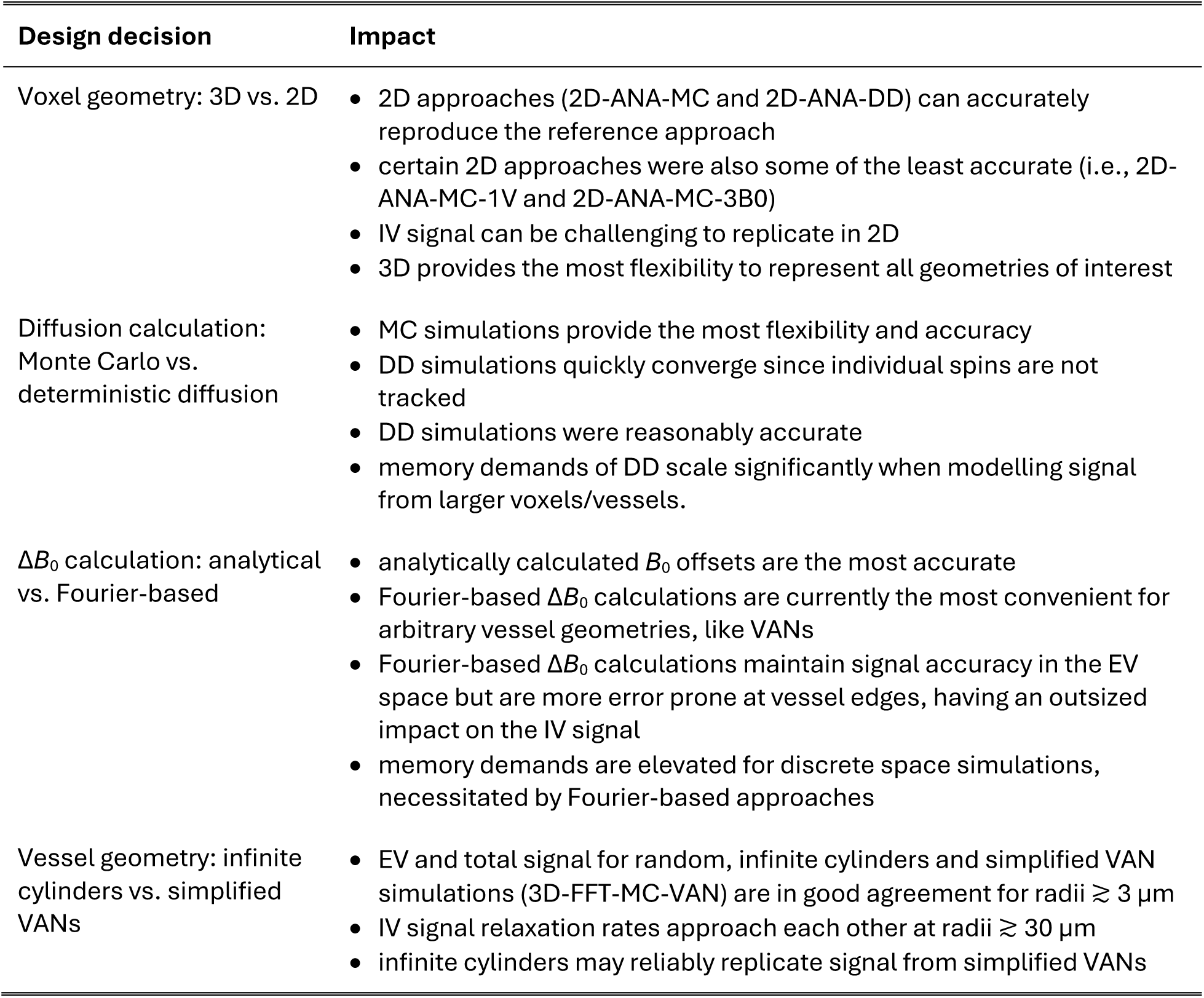
Summary of the impact of the major design decisions one must make when selecting a simulation approach. DD: deterministic diffusion; EV: extravascular; IV: intravascular; MC: Monte Carlo; VAN: vascular anatomical network.

| Design decision | Impact |
| --- | --- |
| Voxel geometry: 3D vs. 2D | • 2D approaches (2D-ANA-MC and 2D-ANA-DD) can accurately reproduce the reference approach<br>• certain 2D approaches were also some of the least accurate (i.e., 2D-ANA-MC-1V and 2D-ANA-MC-3B0)<br>• IV signal can be challenging to replicate in 2D<br>• 3D provides the most flexibility to represent all geometries of interest |
| Diffusion calculation: Monte Carlo vs. deterministic diffusion | • MC simulations provide the most flexibility and accuracy<br>• DD simulations quickly converge since individual spins are not tracked<br>• DD simulations were reasonably accurate<br>• memory demands of DD scale significantly when modelling signal from larger voxels/vessels. |
| $\Delta B_0$ calculation: analytical vs. Fourier-based | • analytically calculated $B_0$ offsets are the most accurate<br>• Fourier-based $\Delta B_0$ calculations are currently the most convenient for arbitrary vessel geometries, like VANs<br>• Fourier-based $\Delta B_0$ calculations maintain signal accuracy in the EV space but are more error prone at vessel edges, having an outsized impact on the IV signal<br>• memory demands are elevated for discrete space simulations, necessitated by Fourier-based approaches |
| Vessel geometry: infinite cylinders vs. simplified VANs | • EV and total signal for random, infinite cylinders and simplified VAN simulations (3D-FFT-MC-VAN) are in good agreement for radii $\gtrsim 3 \mu\text{m}$<br>• IV signal relaxation rates approach each other at radii $\gtrsim 30 \mu\text{m}$<br>• infinite cylinders may reliably replicate signal from simplified VANs |

#### 5.1.1 The impact of using 3D vs. 2D infinite cylinder geometries

The reference technique, 3D-ANA-MC, is based directly on the seminal work of Boxerman, Hamberg, et al. (1995). The method that best agreed with it (in terms of NRMSD and relaxation rates) was, surprisingly, not another 3D method, but rather 2D-ANA-MC (Fig. 6–Fig. 9). Thus, we conclude that 3D field offsets can be well-represented in 2D, likely due to the invariance of the *B*_0_ offsets parallel to an infinite cylinder. Moreover, by assigning a random *B*_0_ direction to each 2D vessel (Fig. 2B), the Δ*B*_0_ distribution of the reference technique was closely replicated (Fig. 10). However, the IV spin-echo Δ*R*_2_ rates were significantly underestimated at small radii using the 2D-ANA-MC method (Fig. 8A.4). This can be explained through imperfect spin refocusing, as follows. Generally, the reason that Δ*R*_2_ is non-zero is that spins cannot be refocused when they sample a different set of Δ*B*_0_ values in the time TE/2 before and after the refocusing pulse. Non-intuitively, for IV spins, since the IV field offset from an infinite cylinder is a constant and would always be refocused, it is actually the *EV* field offsets from neighbouring vessels that give rise to imperfect IV refocusing. For small vessel radii (≲ 10 μm in our simulations), the diffusion length is large relative to the vessel size such that IV spins can sample a wider range of EV field offsets, reducing their likelihood of being refocused. While this is the case for both 2D and 3D geometries, the IV spins in 3D space can diffuse both within the vessel cross-section and along its length (volume *πR*^2^*L*), allowing the spins to sample a wider range of EV fields, whereas IV spins in 2D space are confined to each vessel’s 2D cross-section (area *πR*^2^), limiting the range of Δ*B*_0_ they sample and allowing them to be more readily refocused. Despite this systematic bias, the NRMSD was the smallest for 2D-ANA-MC (Fig. 6 and Fig. 7), translating to perfectly accurate vascular fingerprinting results (Fig. 4 and Fig. 5). Nonetheless, we recognize that more complex geometries than infinite cylinders may not simplify to 2D representations as easily nor produce similarly accurate simulations.

#### 5.1.2 The impact of using different diffusion calculations

The 2D-ANA-DD simulation approach produced excellent agreement with the reference 3D-ANA-MC method, second only to 2D-ANA-MC for the 2D methods. The required spatial resolution was a major consideration when implementing the diffusion calculations with the Monte Carlo or deterministic diffusion approaches, particularly given how changing vessel radii was implemented in this study by scaling the voxel size. The effect of changing the voxel size on the spatial resolution needed to be considered in terms of the resolution of the Δ*B*_0_ maps and diffusion sampling. For the Monte Carlo simulations with Δ*B*_0_ calculated in continuous space (see Table 2), scaling the voxel size had no impact on the resolution of Δ*B*_0_ or diffusion sampling. For the gridded Monte Carlo approaches, the Δ*B*_0_ spatial resolution remained constant, as there were always 6.7 (or 8) grid elements per vessel diameter in 3D (or 2D). However, the diffusion sampling resolution decreased as the voxel size increased. The largest vessel size constrained our resolution since this is where diffusion approaches the static dephasing regime, such that if the resolution was too coarse, spins may never leave the subvoxel they started in. Our grid size convergence simulations ensured that our grid size was adequate in this limiting case (see Supplementary Materials Fig. S1).

For deterministic diffusion, the opposite sampling constraint was true: we previously found that adequately sampling the diffusion kernel was a limiting factor for the accuracy of the simulations (Berman, 2012). As a result, as the vessel radius and voxel size increased, we needed to maintain a minimum spatial resolution such that the diffusion kernel did not become a delta function. Therefore, there were more grid elements per vessel radius as the voxel size increased, such that the effective Δ*B*_0_ resolution increased, but the diffusion sampling remained constant. The results of 2D-ANA-DD may have been improved if even higher spatial resolution were used (see below). However, one concern for the DD method was the increased number of grid elements required when modelling vessels with large radii. Using convolution-based deterministic diffusion, the main advantages of DD over MC methods are that the simulated signals do not have random statistical noise (from an insufficient number of simulated spins), and the computing time is lower since random walks for 10^4^–10^6^ spins are unnecessary. However, as the grid size needs to be increased to adequately sample the diffusion kernel for larger vessel radii, the increased memory demands and convolutions of larger arrays can negate the computational gains relative to the MC computations (Chaussé et al., 2025).

Another limitation of the DD method is the permeability correction, which retrospectively reverses the blurring of magnetization across tissue boundaries (Pannetier et al., 2014). In our experience, when modelling impermeable vessels with DD, results may be erroneous when the diffusion in one timestep allows a significant portion of magnetization to spread diametrically from one side of a vessel to the other. This can be addressed by reducing the timestep (Pannetier et al., 2014) but with the cost of limiting the number of grid points sampling the diffusion kernel, thus necessitating an increased spatial resolution. Furthermore, to the best of our knowledge, only fully permeable or impermeable boundaries have been developed for DD, but not intermediate permeability, which is more easily controlled with MC simulations (Boxerman, Hamberg, et al., 1995). Intermediate permeability could potentially be implemented in the DD framework using an approach developed by Pannetier et al. (2013) for modelling contrast agent extravasation, although this has not been tested.

#### 5.1.3 The effect of using analytical vs. Fourier-based Δ*B*_0_ calculations

As mentioned previously, the interpretation of differences resulting from analytical versus Fourier-based ΔB_0_ calculations, such as 3D-ANA-MC vs. 3D-FFT-MC, are confounded by the fact that the Fourier-based approaches require spatial discretization of the vessels and Δ*B*_0_ distributions. Therefore, the 3D-ANA-MC-GRID simulations were employed to bridge 3D-ANA-MC and 3D-FFT-MC. This spatial discretization introduced only low levels of errors in 3D-ANA-MC-GRID that were relatively constant across radii in the NRMSD (Fig. 6 and Fig. 7). Errors in the Boxerman plots were also small, with the greatest errors observed in the IV Δ*R*_2_ curve at small radii (≲4 μm) (Fig. 9A.4). These discretization errors would have also been present in the DD simulations, and indeed, a similar increase in the IV Δ*R*_2_ values at small radii was observed in 2D-ANA-DD, as opposed to the corresponding *decrease* IV Δ*R*_2_ observed for the non-gridded 2D-ANA-MC simulations (Fig. 8A.4). Since the 3D-ANA-MC-GRID simulations should approach the reference simulations when higher spatial resolutions are employed, the most plausible explanation for the errors at small radii is that the resolution of all these gridded techniques was not entirely adequate.

When using the Fourier transform in discrete space to calculate Δ*B*_0_, 3D-FFT-MC was in excellent agreement with 3D-ANA-MC-GRID and the reference for both the total and EV signals. The Boxerman plots (Fig. 9A.1), which all agreed well, showed nearly identical trends as those from Martindale et al. (2008), who also compared analytical and Fourier-based Δ*B*_0_ calculations using infinite cylinders. Conversely, the Fourier technique dramatically changed the IV Δ*R*_2_ curve relative to discretization alone (3D-ANA-MC-GRID), which, for small radii, could be the result of cancellation of errors from the discretization and the FFT (Fig. 9A.4). The IV Δ*R*_2_ curve appears to have shifted to larger radii (see Fig. 9A.4), but the remaining EV and IV relaxation rates remained consistent with the reference. If there was a corresponding shift to larger radii for the IV Δ*R*_2_*′* curve (Fig. 9A.3), we may not have observed it since the entire curve is already relatively flat across radii. The change in the IV signal, introduced by the Fourier-based Δ*B*_0_ calculation, is a probable source of the outliers in the fingerprinting results at large radii for the 3D-FFT-MC and 3D-FFT-MC-VAN simulations (Fig. 4). At these radii, the *R*^2^ values used for dictionary matching were all very high (>0.99), even across neighbouring radii, such that subtle systematic differences coming from the Δ*B*_0_ calculation could result in erroneous matches. See the next section for discussion on the outlier at 2-μm radius in 3D-FFT-MC-VAN. In general, Fourier-based Δ*B*_0_ calculations are known to be less accurate around discontinuities in susceptibility distributions, such as at vessel walls (Y. C. Cheng et al., 2009; Marques & Bowtell, 2005; Pathak et al., 2008). Since the inaccurate voxels occupy a much larger fraction of the IV space than the EV space, errors are more likely to accumulate in the IV space, consistent with our observations. Overall, however, introducing the Fourier-based Δ*B*_0_ calculation had a minor impact on the NRMSD relative to discretization and on the vascular fingerprinting results.

#### 5.1.4 Infinite cylinders can reproduce simplified VAN simulations

The comparison of infinite cylinders to synthetic VANs seeks to quantify the effect of branching, bending, etc., that are not found in infinite cylinders. The 3D-FFT-MC simulations are useful for interpreting why the synthetic VAN simulations may have differed from infinite cylinder simulations since 3D-FFT-MC and 3D-FFT-MC-VAN are identical in every regard except for the vascular geometry. With the previous Fourier-induced errors in mind, we observed a very different shape of the GE and SE IV Boxerman curves when using the synthetic VANs, while the EV relaxation rates were still relatively consistent with the reference (Fig. 9B). For large radii (≳30 μm), all the relaxation rates converged with those of the 3D-FFT-MC simulations. This convergence is consistent with the fact that the Fourier-based Δ*B*_0_ histograms for infinite cylinders and the synthetic VANs were in close agreement (Fig. 11C) since diffusion effects have a smaller role for larger radii, i.e., in the static dephasing regime (Yablonskiy & Haacke, 1994). The differences in the EV and IV relaxation rates for the VANs at smaller radii (≲30 μm) must, therefore, reflect differences in the specific *history* of field offsets that collections of spins sample in and around branching, tortuous vessels compared to non-branching, infinite cylinders (Kiselev & Novikov, 2018). This is consistent with the findings of Marques & Bowtell (2008), who noted that the inability of infinite cylinders to fully recreate the field offsets at the junctions of branching vessels influences the apparent relaxation rates. These morphologically driven differences in relaxation may also help explain the outlier in the 3D-FFT-MC-VAN fingerprinting results in Fig. 4 at the small radius (2 μm), as there was no corresponding outlier for the FFT-based, infinite cylinder simulations (3D-FFT-MC).

Despite differences in the relaxation rates from the synthetic VANs, the NRMSD of the *total* signals are relatively low for radii ≳ 3 μm (Fig. 6). For the radius of typical capillaries, ∼4 μm, the EV and total NRMSD of the VANs roughly reached a lower plateau (∼7%), suggesting that random infinite cylinders should be a reasonable substitute for modelling signal from the capillary bed. As mentioned in the Methods, we needed to adapt the VAN synthesis to produce the desired CBV with a fixed vessel radius. This resulted in a VAN with 2.25× the normal vessel segment density. Since the field offsets at vessel branching points are poorly captured by infinite cylinders, we expect simulations from the reference approach would have better agreement with simulations from a VAN with a vessel segment density in the normal range since there would be fewer branch points.

Previous work from Marques & Bowtell (2008) also compared simulations from VANs and infinite cylinders. Unlike our results, where the relaxation rates converged at larger radii (Fig. 9B), they observed a more constant, yet slight, overestimation of the VAN relaxation rates in their Boxerman plots across radii. They attributed most of the discrepancy to mismatches in the volume fractions occupied by their VANs vs. infinite cylinders, and they noted that the remaining discrepancies likely came from differences in the field offsets at the junctions of branching vessels. Directly comparing our results to theirs is challenging since they used a finite-difference approach in real space to solve the Bloch equations instead of MC, and a single-vessel model for the infinite cylinder simulations, like our 2D-ANA-MC-1V approach, which we have shown here results in overestimated EV relaxation rates compared to the reference approach. Moreover, they considered a single VAN realization with approximately the same voxel side length as ours (150 μm) but with a distribution of larger radii (2–4.5 μm compared to 1 μm in ours), resulting in far fewer vessel segments, which impacts the ability to produce enough randomness in the distribution of vessel orientations (this was partially compensated by averaging the simulations from multiple *B*_0_ orientations). Pathak et al. (2008) compared VAN-like and infinite cylinder simulations at a single radius of 1.8 μm and found some discrepancies between the simulated time series, similar to what we found at this low radius.

Our synthetic VAN models did not include all aspects of realistic vasculature, nor were they meant to represent vessels such as pial and intracortical vessels, which is why we also referred to them as simplified VANs. The close agreement between our simplified VAN simulations and the other approaches is likely higher compared to what a full VAN or real vasculature might produce because the other aspects of the vasculature (e.g., orientation asymmetries) were absent. Scaling the voxel size to achieve larger vessel sizes, as was done here and is commonly done, does not accurately capture the anatomical properties of the larger vessels. For example, the largest radius simulated here—60 μm—is more representative of pial vessels, which have different branching patterns from the capillary network that we synthesized (Hartung, Badr, Moeini, et al., 2021; Linninger et al., 2019). To achieve a more realistic vascular model using infinite cylinders, these more sparse and structured large vessels, which impose known features on the BOLD signal, such as a cortical orientation dependence (Gagnon et al., 2015; Viessmann et al., 2019), could potentially be modelled through the addition of infinite cylinders with the appropriate size and orientation in place of a full VAN model (Berman, Wang, et al., 2021; Goerke et al., 2007). Although our simulations used a single vessel radius, we would not expect the introduction of vessel-size heterogeneity to impact the performance of the simulation approaches all that much. We would expect that some of the parameters that we examined (e.g., relaxation rates) would become a weighted combination of the results from the individual radii, where that weighting would be sequence specific, as previously demonstrated (Boxerman, Hamberg, et al., 1995; Martindale et al., 2008).

Besides anatomical accuracy, VANs permit realistic flow and oxygen distribution calculations over the continuously connected vascular tree. This continuous distribution of oxygenation levels is harder to reproduce with randomly oriented infinite cylinders; however, it has recently been shown that VANs whose vessels have been compartmentalized into just four oxygenation levels (arteries, capillaries, and two venous compartments) can produce accurate simulations relative to VANs with continuously varying oxygenation levels (Charest et al., 2024). Thus, in the absence of a real, complete VAN, accurate VAN-like simulations may be possible by combining smaller, randomly oriented cylinders, to represent the capillary bed, and some larger, non-randomly oriented cylinders, to represent the remaining vascular compartments, and a small number of oxygenation levels appropriately assigned across the cylinders.

#### 5.1.5 The costs of simplification

While simplifications to the simulation approach reduce computational complexity and promote adoption of forward modelling, there are limits to the extent of simplification. In this regard, the least accurate of all simulations was 2D-ANA-MC-3B0, a 2D approach where the field offsets from two orthogonal orientations were averaged. The error was primarily driven by the IV signal since the distribution of Δ*B*_0_ in the IV space exhibited a localized peak that is not characteristic of random vessel orientations (Fig. 10). Although the blood volume was only 2%, the reference approach’s IV signal accounted for 25% of the total signal’s GE *R*_2_*′* decay (inferred from Fig. 8). Therefore, since 2D-ANA-MC-3B0 was unable to accurately model the IV component’s *R*_2_*′* decay, it is unsurprising that the total signal deviated substantially from the reference, impacting all analyses, including the vascular fingerprinting results.

Notably, the 3*B*_0_ approach is combined with Fourier-based Δ*B*_0_ calculations and deterministic diffusion in another BOLD simulation toolkit, *MrVox2D* (Pannetier et al., 2013). This toolkit was used in the first vascular fingerprinting study (Christen et al., 2014) and several other studies that followed. However, as we have shown here, the 3*B*_0_ approach, Fourier-based Δ*B*_0_ calculations, and DD are all prone to errors that can lead to biased fingerprinting results (Fig. 4 and Fig. 5). Consistent with our observed overestimation of the known vascular radii, the contributors to *MrVox2D* have recently shown that the 3*B*_0_ approach overestimated the underlying distribution of radii by 14–19%, on average, as compared to a more realistic VAN-based dictionary when performing vascular fingerprinting on *in vivo* rat data (Delphin et al., 2024). Thus, the 3*B*_0_ approach should be used with caution.

Another simplification we tested was the single-vessel approach, 2D-ANA-MC-1V. An advantage of this approach over others is that a small voxel size is required since only a single vessel must be contained within it. This can translate to fewer MC spins to be simulated or a smaller grid size for DD simulations, albeit at the expense of having to iterate at varying polar *B*_0_ angles. However, this single-vessel simplification showed poorer agreement with the reference method than the other 2D approaches (save for the 3*B*_0_ method) as it produced larger NRMSDs (Fig. 7), increased EV relaxation rates, and decreased IV relaxation rates (Fig. 8B). The elevated EV relaxation rates could be the result of inadequately sampling the distribution of polar angles from 0 to 90°, since Mueller-Bierl et al. (2007) showed that the EV relaxation rates are overestimated when using 6 or 9 polar angles compared to when using 18 (our study used 9 angles, like Uludag et al. (2009)). The underestimation of the IV relaxation rates could be mitigated by scaling the IV signal by the EV signal, as proposed by Boxerman, Bandettini, et al. (1995), since the IV spins should sense the EV field from neighbouring vessels. Despite these biases, the single-vessel approach has been used in seminal works that have helped establish our biophysical understanding of the BOLD signal, such as Ogawa et al. (1993) and Uludag et al. (2009), using 16 and 9 polar angles, respectively. If one uses their corresponding results to consider *relative* trends that relate the underlying physiology to the BOLD signal (e.g., the microvascular specificity across field strengths) rather than defining specific relationships to later quantify physiological parameters *in vivo* (e.g., quantifying CBV based on relaxation rates), the trade-off between quantitative accuracy and ease of implementation is reasonable.

### 5.2 Limitations and future work

#### 5.2.1 Breadth of simulation approaches vs. simulation parameter values

One limitation of this work is that a single set of simulation parameters (i.e., including susceptibility offset, blood volume, field strength, and pulse sequence) was used. Many previous studies have looked at the impact of these parameters on transverse signal decay using their preferred simulation approach (e.g., Boxerman, Hamberg, et al. (1995), Martindale et al. (2008), Uludag et al. (2009), etc.). However, our emphasis was to explore the mathematical differentiations instead of physiological/acquisition differentiations, and our choice of parameters allowed us to do so. More specifically, the parameter values in Table 1 were chosen to be consistent with typical fMRI studies, i.e., cortical venous blood volume (2%), venous Δ*χ*, field strength (3 T), pulse sequence (TE = 30 ms for GE, TE = 70 ms for SE), allowing for a representative range of signal characteristics to be captured, such as GE decay and SE refocusing. If additional simulation settings were run, we would expect the absolute value of many of the simulation results to change, and perhaps some simulations would perform better than with our original settings; however, we would not expect wholesale differences that would dramatically impact the conclusions drawn. The impact of including other simulation parameters, such as vessel size heterogeneity and continuously varying blood oxygenation—physiological features that VANs excel at modelling—is discussed in more detail below.

This study provides a framework for evaluating other simulation approaches that were not examined here. While we did explore many approaches, there are several more equally valid ones that we did not include, e.g., forcing spins to reflect off of vessel walls rather than reseeding their positions when they cross a boundary (Dickson et al., 2010), or more recent rapid simulation techniques combining histogram-based Δ*B*_0_ calculations and deep learning-based diffusion calculations (Coudert et al., 2025).

#### 5.2.2 Intravascular signal modelling

For all techniques considered, the relative error was greatest for the IV signal. In reality, the IV signal for all approaches was still not physically accurate since uniformly magnetized cylinders were used, rather than explicitly modelling the fields of individual red blood cells. For blood vessels with flows ≳ 20 mm/s, uniform cylinders may accurately capture the IV and EV signal (Boxerman, Hamberg, et al., 1995; Martindale et al., 2008), but more advanced models may be necessary for capturing the IV signal from capillaries with slow blood flow. For partially deoxygenated venous or capillary blood, diffusion and exchange through red blood cells decreases the IV *T*_2_ (Berman & Pike, 2018), which further down-weighs the contribution of the IV signal to the total signal, thus reducing the magnitude of the relaxation errors imparted by the IV signal. For instance, at 3 T, the *T*_2_ of 65% oxygenated blood and of EV tissue are around 50 ms and 75 ms, respectively (Uludag et al., 2009). Thus, if our simulations had considered intrinsic *T*_2_ decay, there would be a slight reduction in the errors in the total (EV+IV) signal, impacting the NRMSD, relaxation rates, and fingerprinting results. At higher field strengths, *T*_2_ is considerably shorter in veins than in EV tissue, so we would expect an even smaller contribution of the IV signal to the NRMSD, therefore, potentially permitting more flexibility in the choice of simulation approach at ultra-high fields (Uludag et al., 2009).

While intrinsic *T*_2_ decay would reduce the relative magnitude of the IV errors in the total signal, the overall error could become much more substantial if CBVs greater than 2% were considered. Large blood vessels, comprising the macrovasculature, can occupy much larger fractions of a voxel than 2% and have a considerable influence on the spatiotemporal characteristics of the fMRI signal, as demonstrated theoretically and empirically (Hyde & Li, 2014; Menon, 2002; Zhong, Tong, et al., 2024). Therefore, studies that attempt to model macrovascular contributions to the BOLD signal may be more sensitive to the specific simulation approach (Zhong, Polimeni, et al., 2024). Modelling non-BOLD sequences that have a greater relative contribution from IV spins could also be more sensitive to these IV-related inaccuracies, e.g., in functional MR angiography (Cho et al., 2012) or arterial spin labelling with background suppression (Germuska et al., 2019). If inadequate spatial resolution was the major source of IV error for some of the approaches (e.g., the Fourier-based approaches), then this could be improved by simply increasing the resolution, since spatial sampling requirements are typically constrained by the *microvasculature*, not the macrovasculature.

#### 5.2.3 Choice of reference simulation approach

Finally, all our analyses compared each simulation approach to random, infinite cylinders using analytically calculated field offsets and Monte Carlo diffusion (3D-ANA-MC). The motivations for using infinite cylinders for the reference included the wealth of literature and experience within the community with using infinite cylinders and the availability of analytical solutions for the field offsets with well-understood signal characteristics. However, we ultimately want our simulations to reflect the signal induced by *realistic* vasculature, as in VANs, which can more readily capture a range of vascular properties, as described above in section 5.1.4. The challenge with VAN-based simulations is that they are currently calculated with spatial discretization and Fourier-based Δ*B*_0_ calculations, which have known inaccuracies, as discussed above. Furthermore, it is unclear how results from VANs generalize to other brain areas since most VANs, including synthetic VANs, are derived from in vivo measurements that are regionally varying. Therefore, VAN-based simulations may give a more regionally biased approximation of signal across the brain, whereas infinite cylinder simulations may be more generalizable, albeit not perfectly representing signal from any single brain area. Building on our results, in future work, we plan to further investigate optimal strategies for implementing the most realistic VAN-based simulations, accounting for the accuracy of the results and the computational demands necessitated by the spatial discretization.

## 6 Conclusions

This study has explored the relative agreement of a wide range of BOLD signal biophysical simulation approaches and compared their computational performance (i.e., memory and run time). All simulations were implemented in the BOLDsωimsuite toolkit. By incrementally varying simulation design choices (see Fig. 3), we were able to isolate the changes introduced by the various approaches. This work was the first to explicitly show the equivalence of the *B*_0_ distributions and the near-perfect equivalence of the simulation results of the 2D simulation approach with random *B*_0_ directions per vessel (2D-ANA-MC) compared to the 3D reference (3D-ANA-MC). While most of the simulations were in good agreement with the reference approach, our implementations of the single-vessel approach (2D-ANA-MC-1V) and the 3*B*_0_ approach (2D-ANA-MC-3B0) both had poorer agreement by most metrics considered. Finally, we showed that infinite cylinders may be a reasonable approximation of the capillary network—as modelled using synthetic VANs—for simulations using vessel radii ≳ 3 μm. We conclude, therefore, that the 3D-ANA-MC reference approach is an attractive option for exploratory simulations, balancing ease of implementation, accessibility, versatility, computational efficiency, accuracy of results, and interpretability.

## 7 Ethics

Not applicable, as no human or animal data were used.

## 8 Data and Code Availability

The BOLDsωimsuite simulation toolkit is openly available at https://github.com/jacobchausse/BOLDswimsuite. The data and analysis code that support the findings of this study are available from the corresponding author upon reasonable request.

## 9 Author Contributions

**Avery J. L. Berman:** Conceptualization, Methodology, Investigation, Formal analysis, Writing – Original Draft, Writing – Reviewing & Editing, Visualization, Funding acquisition. **Jacob Chaussé:** Methodology, Investigation, Software, Writing – Reviewing & Editing. **Grant Hartung:** Software, Resources, Writing – Reviewing & Editing. **Jonathan R. Polimeni:** Conceptualization, Writing – Reviewing & Editing. **J. Jean Chen:** Conceptualization, Resources, Writing – Reviewing & Editing, Funding acquisition, Project administration.

## 11 Acknowledgements

We are grateful to Prof. Bruce Pike for many insightful discussions on BOLD modelling. The simulations in this study were enabled in part by support provided by Compute Ontario (https://www.computeontario.ca/) and the Digital Research Alliance of Canada (alliancecan.ca). Financial support comes from NSERC (RGPIN-2022-04886), CIHR (MFE-164755 and FDN-148398), the Canada Research Chairs Program (JJC), the NIH NIBIB (grants P41-EB030006 and R01-EB032746), the NIH NINDS (grant R01-NS128843), and by the *BRAIN Initiative* (NIH NIMH grants R01-MH111419, R01-MH111438, F32-MH125599, and U19-NS123717).

