## Supplementary Material for "Evaluating BOLD functional MRI biophysical simulation approaches: impact of vascular geometry, magnetic field calculations, and water diffusion models"

### S1 Effective voxel sizes

**Table S1:** Summary of the voxel sizes used across simulation approaches for each vessel radius. The columns provide the vessel radii, effective voxel edge length for each radius for all 3D methods, 2D-ANA-DD, and 2D-ANA-MC-1V, the number of grid elements for all discrete 3D methods and 2D-ANA-DD, and the resulting grid size for the discrete 3D methods and 2D-ANA-DD.

| Vessel radius (μm) | Effective voxel edge length, 3D methods (μm) | Effective voxel edge length, 2D-ANA-DD (μm) | Effective voxel edge length, 2D-ANA-MC-1V (μm) | # grid elements, discrete 3D methods | # grid elements, 2D-ANA-DD | Grid size, $\Delta x$, discrete 3D methods (μm) | Grid size, $\Delta x$, 2D-ANA-DD (μm) |
|---|---|---|---|---|---|---|---|
| 1 | 149 | 251 | 12.5 | $500^3$ | $1000^2$ | 0.298 | 0.251 |
| 1.5 | 223.5 | 376.5 | 18.8 | $500^3$ | $1000^2$ | 0.447 | 0.377 |
| 2 | 298 | 502 | 25.1 | $500^3$ | $1000^2$ | 0.596 | 0.502 |
| 3 | 447 | 753 | 37.6 | $500^3$ | $1000^2$ | 0.894 | 0.753 |
| 4 | 596 | 1004 | 50.1 | $500^3$ | $1060^2$ | 1.192 | 0.947 |
| 5 | 745 | 1255 | 62.7 | $500^3$ | $1330^2$ | 1.490 | 0.944 |
| 6 | 894 | 1506 | 75.2 | $500^3$ | $1590^2$ | 1.788 | 0.947 |
| 8 | 1192 | 2008 | 100.3 | $500^3$ | $2120^2$ | 2.384 | 0.947 |
| 10 | 1490 | 2510 | 125.3 | $500^3$ | $2650^2$ | 2.980 | 0.947 |
| 13 | 1937 | 3263 | 162.9 | $500^3$ | $3440^2$ | 3.874 | 0.949 |
| 16 | 2384 | 4016 | 200.5 | $500^3$ | $4230^2$ | 4.768 | 0.949 |
| 20 | 2980 | 5020 | 250.7 | $500^3$ | $5290^2$ | 5.960 | 0.949 |
| 30 | 4470 | 7530 | 376.0 | $500^3$ | $7930^2$ | 8.940 | 0.950 |
| 40 | 5960 | 10040 | 501.3 | $500^3$ | $10570^2$ | 11.920 | 0.950 |
| 60 | 8940 | 15060 | 752.0 | $500^3$ | $15860^2$ | 17.880 | 0.950 |

### S2 Vascular Anatomical Network (VAN) synthesis and simulation details

#### S2.1 VAN synthesis and binarization

To synthesize the VANs, a large voxel (1 $mm^3$) was required to satisfy the statistical constraints (i.e., tortuosity and branching patterns) of the capillary VAN model (Hartung et al., 2021, 2025; Linninger et al., 2019). To generate voxels for signal simulations with the same characteristics as our infinite cylinder voxels, specifically the ratio of the vessel diameter to the voxel edge width, we extracted 10 sub-voxels from the larger VAN. Any vessel segments that exited the sub-voxels were pruned, resulting in a slightly reduced blood volume than what was nominally set. The nominal blood volume was 2%, and the resulting mean and standard deviation of the extracted VANs were (1.79 ± 0.09)%.

Calculating the $\Delta B_0$ distributions using the Fourier approach requires a discretized susceptibility map. For voxels with a single $\Delta\chi$ across all vessels, as in this study, a discrete susceptibility map can be generated from a vessel mask. The vessel masks were generated by convolving the vessel centrelines with a sphere of the desired radius. In practice, this was achieved by filling the voxel with a binary sphere mask that moved along the centreline of each vessel using a step size of 5% of the radius (i.e., 0.05 μm steps for 1-μm vessels).

#### S2.2 Signal blood volume correction

To remove the effect of the small blood volume difference when comparing simulations, we used the following approach (Stone et al., 2019). The extravascular (EV) signal from a vessel network can, in general, be expressed as $S(t) = \exp[-V \cdot f(t)]$, where $V$ is the blood volume fraction and $f(t)$ is a sequence-dependent function representing the amount of attenuation over time (Kiselev & Posse, 1999). Therefore, the original EV simulated signals from the VANs, $S_{EV}(t)$, were scaled as $S'_{EV}(t) = [S_{EV}(t)]^{V_0/V}$, where $V_0$ = 0.02 and $V$ was the corresponding VAN's blood volume, and the total signal was recalculated as $S'_{tot}(t) = (1 - V_0)S'_{EV}(t) + V_0 S_{IV}(t)$. No change was made to the intravascular signal ($S_{IV}$).

### S3 Methodology to evaluate computational performance

Computational performance across most approaches was evaluated using the same methods described by Chaussé et al. (2025). Briefly, simulations were run on 10 random voxels with 400 infinite cylinders at 2% cerebral blood volume for each approach (except for 2D-ANA-MC-1V, which used a single voxel with a single vessel). All simulations were run for 600 time steps (with 40,000 spins for the MC simulations). The grid size was set to 1000 for 2D-ANA-DD and 500 for 3D-ANA-MC-GRID and 3D-FFT-MC. For each simulation, the computing time and peak memory demand were recorded. The computing time was divided into time to initialize the simulations (e.g., create the set of vessels or, if necessary, calculate the $\Delta B_0$ maps) and the run time to calculate the MR signal. Note, the performance of 2D-ANA-MC-3B0 is comparable to 2D-ANA-MC, therefore, it was not evaluated. Additionally, preparing the VAN vessel masks required several steps that were not evaluated for the computing requirements; therefore, the results for 3D-FFT-MC-VAN are not shown. After preparing the vessel masks, the simulations would be comparable to 3D-FFT-MC. The computational time and memory requirements for the 2D-ANA-DD simulations increased nearly exponentially with vessel radius, as reported by Chaussé et al. (2025), and only the minimum requirements for a 1-μm vessel radius were considered here.

Unlike the simulations for the accuracy comparisons in the main article, these simulations ran on a different computing cluster with AMD EPYC 7532 (Zen 2) 2.40-Gz CPUs.

### S4 Results

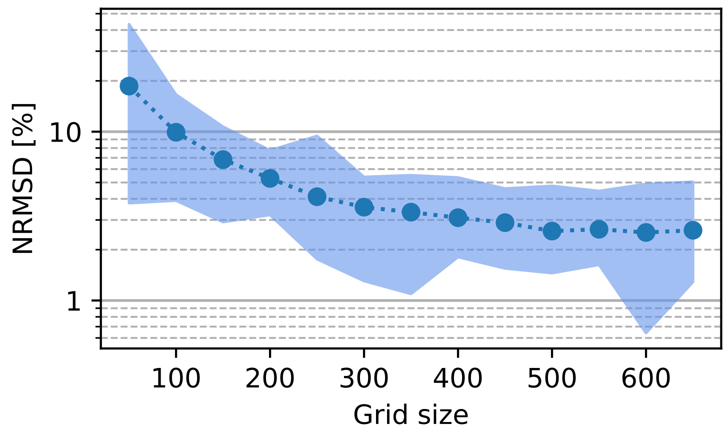


**Fig. S1:** Convergence of the total signal (EV+IV) of the 3D gridded, analytical $\Delta B_0$, Monte Carlo approach (3D-ANA-MC-GRID) as a function of the grid size along each dimension. Convergence was assessed using the normalized root mean square difference (NRMSD) relative to the reference approach (3D-ANA-MC), as described by Eq. [13] in the main text. The simulated signal evolutions from ten voxels were averaged before calculating the NRMSD. The circle markers represent the mean NRMSD across all radii, and the shaded region represents the minimum and maximum NRMSE over all radii. The mean NRMSD plateaued at a grid size of 500.

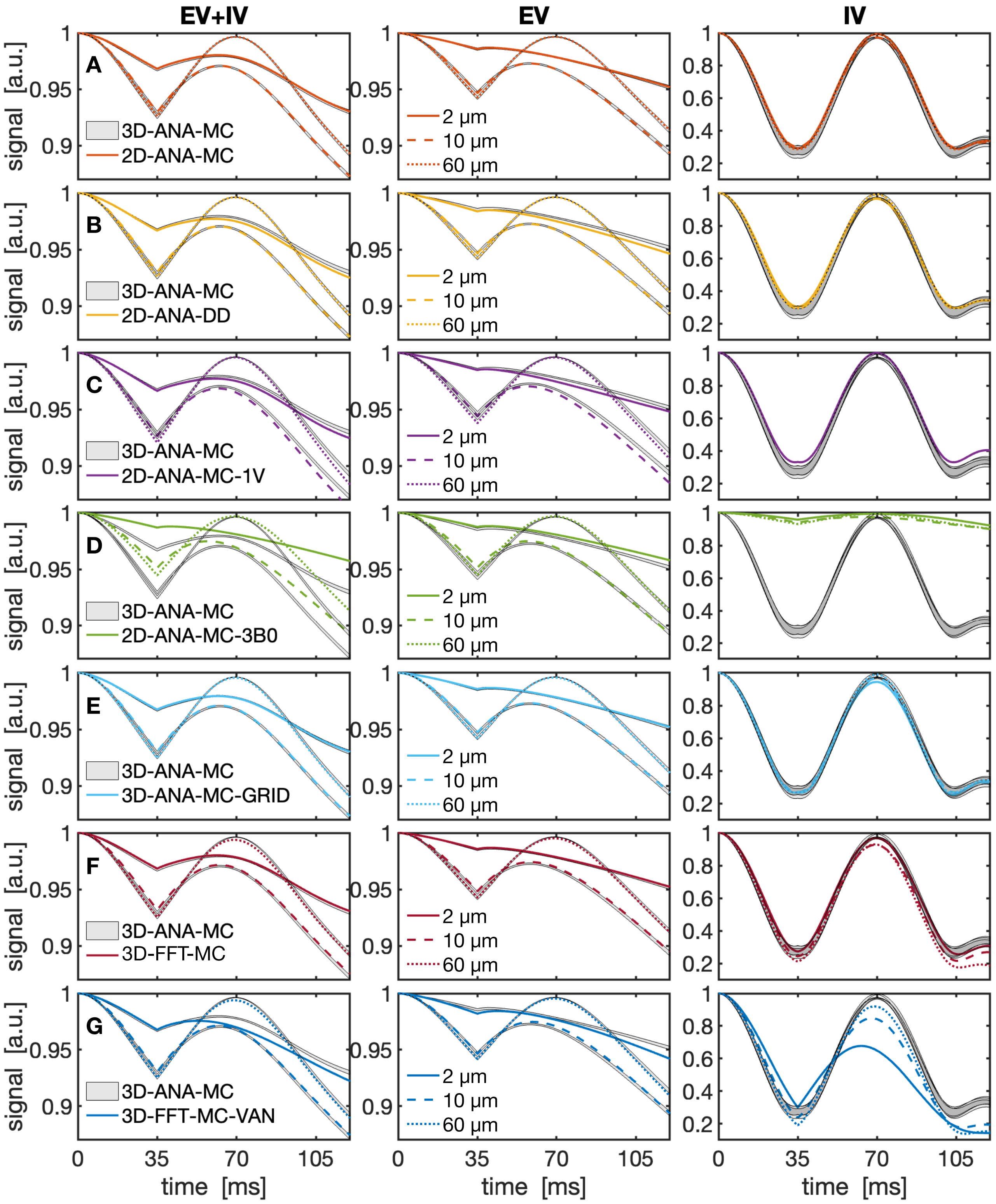


**Fig. S2:** Mean simulated spin-echo signal evolution magnitude (TE = 70 ms) for each simulation technique at three representative radii. The total signal (EV+IV), EV only, and IV only are plotted across columns, each normalized to 1 at time point 0. The grey shaded bars in all plots represent the mean reference simulation (3D-ANA-MC) plus/minus one standard error over all ten voxels. (A–G) The mean time series from the remaining simulation approaches are plotted with the coloured lines, with each radius designated by the line style; error bars are omitted for clarity.

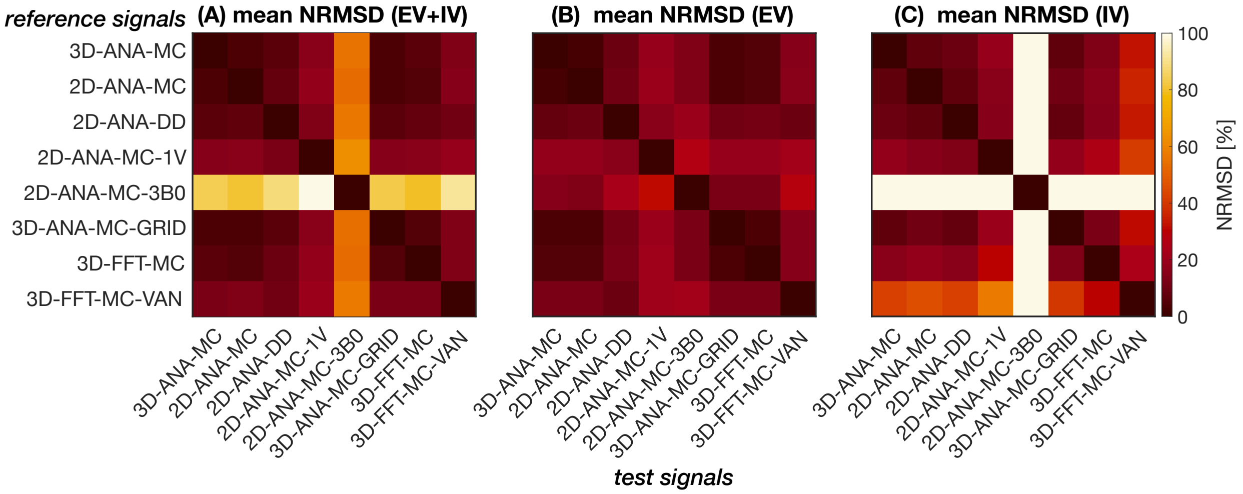


**Fig. S3:** The normalized RMS difference (NRMSD) averaged over all radii for each simulation technique (columns) relative to the reference signals (rows). NRMSD is plotted for (A) the total signal (EV+IV), (B) EV only, and (C) IV only. All plots share the same colour bar as (C).

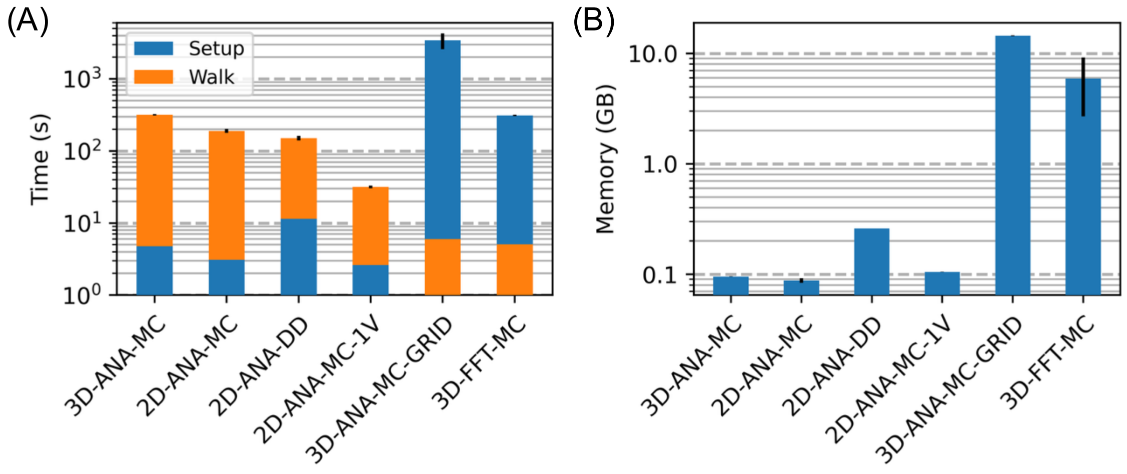


**Fig. S4:** Computational times and memory requirements from sample simulations. (A) Computational times, divided into voxel generation time ("Setup") and diffusion and signal calculation time ("Walk"), with the lower of the two times displayed on the bottom portion of the bar due to the logarithmic y-axis. (B) Peak memory requirement during simulation time. Note, the performance of 2D-ANA-MC-3B0 is comparable to 2D-ANA-MC, therefore, their results are not shown. Additionally, preparing the VAN vessel masks required several steps that were not evaluated for the computing requirements; therefore, the results for 3D-FFT-MC-VAN are not shown. After preparing the vessel masks for the VAN, the simulations would be comparable to 3D-FFT-MC. The computational time and memory requirements for the DD simulations increased nearly exponentially with vessel radius, as reported by Chaussé et al. (2025), and only the minimum requirements are plotted above for a 1-μm vessel radius.
